# Single-stranded DNA binding proteins are essential components of the architectural LDB1 protein complex

**DOI:** 10.1101/2025.06.05.658047

**Authors:** Xiaokang Wang, Nicholas G. Aboreden, Ying Cai, Jessica C. Lam, Kate A. Henderson, Jiaqi Xiang, Belinda M. Giardine, Ross C. Hardison, Cheryl A. Keller, Lalitha Nagarajan, Stephen J. Brandt, Gerd A. Blobel

## Abstract

Transcriptional enhancers are brought into proximity with promoters via chromatin looping. The architectural transcription cofactor LDB1 facilitates spatial connectivity among enhancers and promoters but whether this occurs through simple dimerization or requires partner molecules is unknown. Here we investigated single-stranded DNA binding proteins (SSBPs), known LDB1 interactors, in regulating LDB1-mediated chromatin looping and transcription. SSBP2, SSBP3, and SSBP4 colocalize with LDB1 genome wide. Among these, only SSBP3 is essential for erythroid cell viability, LDB1 function, and transcription. LDB1, but not single-stranded DNA, is the predominant genome-wide tether of SSBP3 to chromatin. Notably, SSBP3 depletion for under one hour in SSBP2/4 knockout cells globally weakened LDB1-dependent chromatin loops and lowered nascent transcription without impacting LDB1’s chromatin binding. Chromatin tethering experiments revealed SSBP3 and LDB1 mutually depend on each other to form looped contacts. SSBP3 stabilizes LDB1 homodimers in solution providing a possible mechanism of action. In sum, SSBPs emerge as key functional components of the architectural LDB1 complex, shedding new light on the regulation of enhancer-promoter interactions and gene expression.

## Introduction

Enhancers specify time, place and level of gene transcription^1,2^. How enhancers communicate with their target promoters has been a major focus in the field of gene regulation. Several models have been proposed to explain the mechanisms by which enhancers communicate with promoters, including chromatin looping, which is supported by substantial evidence^1–4^. How looped enhancer-promoter (E-P) contacts are established mechanistically, and how specificity is achieved are subjects of intense investigation. In one model, transcription factors recruit co-factors that facilitate long range contacts, perhaps via the formation of dimers or multimers. One such co-factor is LDB1, which plays a role in E-P looping in various cellular contexts^5^ (For review see [^6–10^])

LDB1 is widely expressed and does not bind DNA directly. It is directed to its targets by gene specific transcription factors such as GATA1 and TAL1, with LMO2 acting as a bridge^11^ (For review see [^8,12,13^]). In other contexts, LDB1 recruitment is mediated by LIM homeodomain proteins, such as LHX1^14^ (For review see [^9,10,15–17^]).

Knockdown of LDB1 in erythroid cells reduced the formation of a chromatin loop between the β*-globin* gene promoter and its upstream enhancer, called the Locus Control Region (LCR), and diminished β*-globin* transcription^5,18,19^.

Gain-of-function experiments cemented a direct role for LDB1 in chromatin looping and supported the idea that chromatin loops can cause and not just be a consequence of enhancer driven transcription^20–22^. Specifically, artificial tethering of LDB1 to the β*-globin* promoter induced E-P looping and activated transcription^20,21^. Forced LDB1 recruitment also enabled E-P rewiring to transcriptionally re-program the human β*-globin* locus^21,23,24^.

On a broader scale, LDB1 is required for E-P loops at a substantial fraction of genes^25^. Such LDB1-dependent loops include long range contacts that can even reach across boundaries of topologically associating domains (TADs). Importantly, most LDB1 anchored loops, including some spanning up to a megabase, can occur in the absence of the cohesin/CTCF machinery, suggesting that LDB1 can form strong contacts independently of chromatin loop extrusion, presumably via protein-protein interactions^25^. Moreover, in murine olfactory sensory neurons LDB1 can organize inter-chromosomal enhancer hubs to activate olfactory receptor gene transcription^26^.

The molecular basis for how LDB1 forms contacts *in vivo* is still unclear. The LDB1 N-terminus contains a dimerization domain (DD)^27–29^ which contributes to its looping function (Figure 1)^20–22,24^. In addition, LDB1 harbors an LDB/Chip conserved domain (LCCD)^30^ which has been proposed to interact with CTCF to mediate heterotypic LDB1-CTCF anchored loops^31^. LDB1 also interacts with a class of molecules termed single-stranded DNA-binding proteins (SSBPs), also known as sequence-specific single-stranded DNA-binding proteins (SSDPs)^32^ (For review see [^10^]).

**Figure 1.**
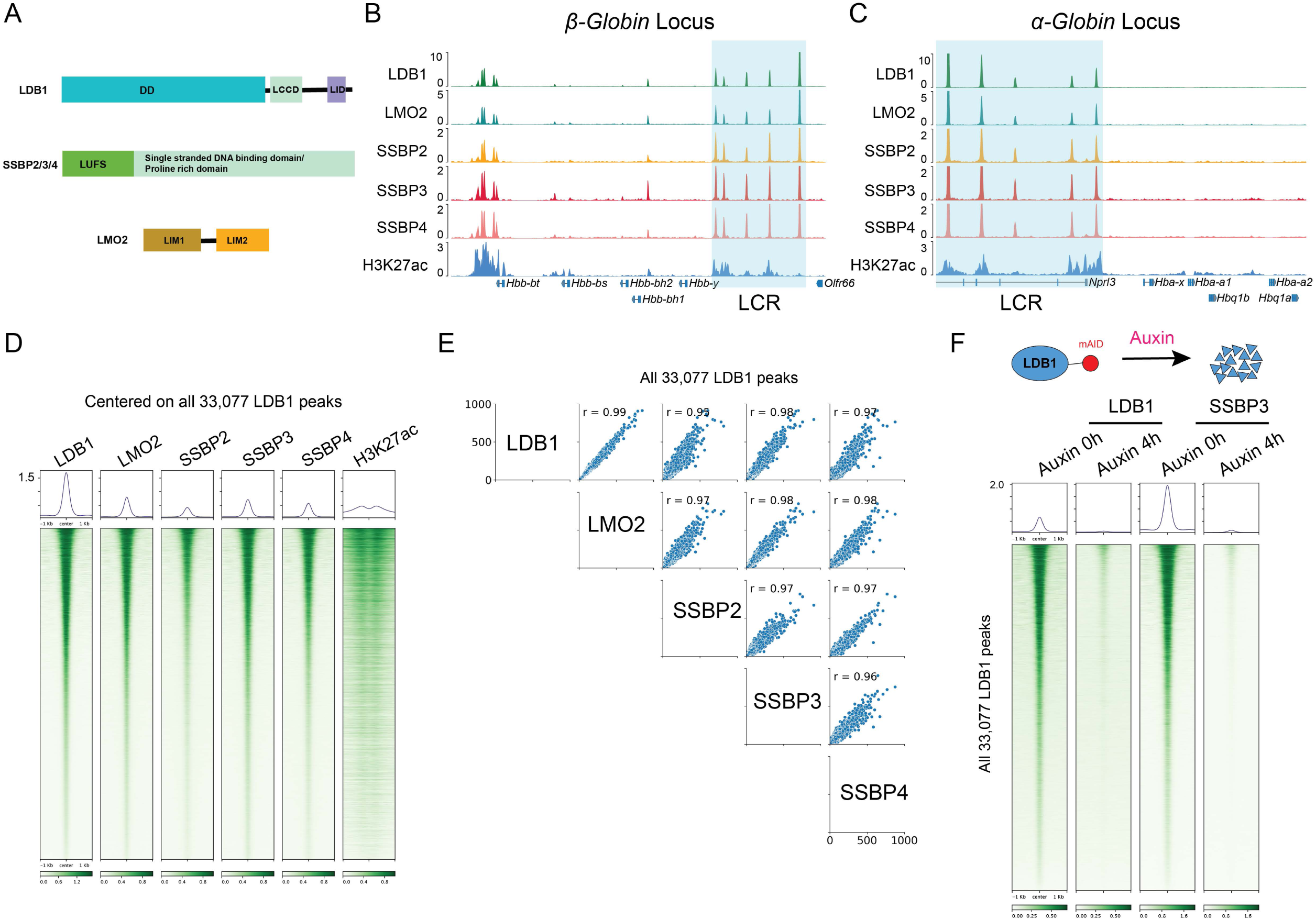
SSBP2/3/4 colocalize with LDB1 genome-wide. (A) Domain organization of LDB1, SSBP2/3/4, and LMO2 (B-C) ChIP-seq tracks showing colocalization of LDB1 (n=5), LMO2 (n=3), SSBP2 (n=5), SSBP3 (n=2), SSBP4 (n=4), and H3K27ac (n=2) at the β*-globin* locus (B) and α*-globin* locus (C). LCR is highlighted in cyan. (D) Heatmaps centered on LDB1 peaks showing a genome wide colocalization of LDB1 (n=5), LMO2 (n=3), SSBP2 (n=5), SSBP3 (n=2), SSBP4 (n=4), and H3K27ac (n=2). (E) Pairwise scatter plots and pairwise Pearson correlation coefficients among peak strengths of LDB1 (n=5), LMO2 (n=3), SSBP2 (n=5), SSBP3 (n=2), and SSBP4 (n=4). (F) Upper: diagram showing that LDB1 was depleted by auxin treatment. Lower: heatmaps showing LDB1 (n=2) and SSBP3 (n=2) chromatin binding before and after 4h auxin treatment in LDB1-AID cells.

The first SSBP was isolated in chickens through a screen for nuclear factors that bind single-stranded DNA (ssDNA) in a sequence specific manner^33^. Subsequently, SSBP2, SSBP3 (also known as SSDP1), and SSBP4, were identified in mammals^34,35^.

SSBPs are involved in multiple biological processes, and misexpression or mutations are associated with some neurodevelopmental disorders and cancers. SSBP3 regulates glucose homeostasis, pancreatic islet architecture, and beta-cell identity^36^. Alterations in *SSBP3* gene expression have been associated with autism^37^, and mutations of *SSBP3* can cause a headless phenotype in mouse embryos^38,39^. Several chromosomal translocations which result in SSBP2 fusion proteins have been reported in B-cell acute lymphoblastic leukemia (B-ALL)^40–42^.

SSBPs were found to be components of LDB1-associated complexes by affinity purification using LDB1 as bait in HeLa cells^32^, and LDB1-SSBP interactions were subsequently found in other cell types and organisms^30,32,43–49^. Although SSBPs can bind ssDNA^33^ in gel-shift assays, to our knowledge there is little evidence that SSBPs bind ssDNA *in vivo*.

Human and mouse genomes encode three SSBPs: SSBP2, SSBP3, and SSBP4, which have high sequence similarity (Figure S1A) while SSBP1 is unrelated. During human and mouse hematopoiesis, SSBP3, alongside other erythroid transcription factors and cofactors, including LDB1, is highly expressed in erythroid cells, while SSBP2 and SSBP4 are mainly expressed in immune cells (Figure S1B and C)^50,51^.

SSBPs have an N-terminal LUFS (LUG/LUH, Flo8, and SSBP/SSDP) domain^30^ and a C-terminal single stranded DNA binding domain (also called proline rich domain)^33,39^ (Figure 1A). Structural studies revealed the conformation of the LDB1/SSBPs complex^47,52–54^.

SSBPs were suggested to stabilize LDB1 by preventing its RLIM-mediated ubiquitination and proteasomal degradation^55,56^. However, these findings were based on long-term overexpression or depletion experiments^55,56^, which can confound interpretation. One study argued that SSBPs do not regulate LDB1 protein levels, and that, conversely, LDB1, which has a longer half-life (∼27 hours) than SSBPs (5 to 8 hours), enforces stability on SSBPs^44^. Generally, it is quite common that removing one component of a given protein complex can destabilize others, which does not necessarily mean that maintaining protein stability is its principal function. SSBPs may also contribute to LDB1 function via other mechanisms, such as promoting LDB1 dimerization or oligomerization. Indeed, previous studies suggested that LDB1 exhibits weak affinity for itself ^27,29^, raising questions as to how it could form chromatin loops on its own.

Here, we comprehensively studied the functions of SSBP2/3/4 proteins in G1E-ER4 cells, a *GATA1*-null erythroid cell line expressing GATA1 protein fused to the hormone-binding domain of the mammalian estrogen receptor (GATA1-ER). Treatment with estradiol activates GATA1 and induces erythroid maturation in a manner very similar to that of primary erythroid cells^57^. We found that SSBP2/3/4 colocalize with LDB1 genome-wide in both immature and mature erythroid cells. LDB1 is the predominant, if not only, tether that recruits SSBP3 to chromatin. *In vitro*, the dimerization of LDB1 is significantly enhanced by SSBP3. Depletion of SSBP3 for under one hour does not affect LDB1 stability on chromatin, contrasting with previous reports that employed prolonged perturbations^55,56,58–60^. Instead, our results revealed that SSBP3 directly supports LDB1’s role in E-P looping and transcription activation. Our findings nominate SSBP3 as an essential component of the architectural LDB1 complex. We propose that depending on tissue distribution, other SSBP proteins function similarly to establish specific long range regulatory contacts in the context of LDB1.

## Results

### SSBP2/3/4 colocalizes with LDB1 genome-wide

To examine whether SSBPs bind to chromatin and if so, whether they colocalize with LDB1, we performed chromatin immunoprecipitation followed by sequencing (ChIP-seq) for SSBP2/3/4 and LDB1 in undifferentiated G1E-ER4 cells.

We first optimized the ChIP protocol by comparing two crosslinking methods: dual crosslinking (disuccinimidyl glutarate plus formaldehyde)^61^ and single crosslinking (formaldehyde). We found that dual crosslinking was essential for SSBP3 ChIP and vastly improved ChIP for LDB1 (Figure S1D and E).

Due to the lack of ChIP grade antibodies against SSBP2 and SSBP4, we knocked-in an eGFP-mAID-2xFLAG tag at the C-terminus of SSBP2 (the mAID was not put into use as SSBP2 is not essential for cell viability) and a 2xFLAG tag at the N-terminus of SSBP4 (rather than the C-terminus as two splicing forms exist that alter the C terminus)(Figure S1F), enabling anti-FLAG ChIP.

We also performed ChIP for LMO2, which links GATA and TAL proteins to LDB1 via LDB1’s LID domain^62,63^ (Figure 1A). In total, we identified 33,077 LDB1 peaks, 18,867 LMO2 peaks, 14,292 SSBP2 peaks, 21,145 SSBP3 peaks, and 15,672 SSBP4 peaks in undifferentiated G1E-ER4 cells.

SSBP2, SSBP3, and SSBP4 strongly colocalized with LDB1, LMO2, and H3K27ac. For example, high signals for binding of each of these proteins and H3K27 acetylation are observed in both the α*-globin* and β*-globin* loci at the enhancer regions (Figure 1B and 1C). A heatmap centered on all 33,077 LDB1 peaks reveals co-enrichment of LMO2, SSBP2/3/4, and H3K27ac (Figure 1D), and binding intensities of LDB1, LMO2, and SSBP2/3/4 are strongly correlated at all LDB1-occupied sites (Pearson correlation coefficients ≥0.95 (Figure 1E)). Genome-wide, 99.42% of LMO2 peaks, 97.03% of SSBP2 peaks, 94.29% of SSBP3 peaks, and 97.32% of SSBP4 peaks overlap with LDB1 peaks (Figure S1G).

In contrast, not all LDB1 peaks are co-occupied by SSBPs (Figure S1G), and we divided the 33,077 LDB1 peaks into two categories: 1) 20,963 LDB1 peaks that overlap with at least one SSBP; and 2) 12,114 LDB1 peaks that do not overlap with any SSBP. At regions where LDB1 overlapped with at least one SSBP, LDB1, LMO2, SSBP2/3/4, and H3K27ac signals were stronger when compared to sites at which LDB1 did not overlap with any SSBPs (Figure S1H). However, at the 12,114 LDB1 peaks classified as not co-occupied by SSBPs, weak enrichment was still visible, indicating that this classification was due to thresholding and/or the use of different antibodies (Figure S1H). Collectively, these results demonstrate that SSBP2/3/4 and LDB1 are highly colocalized genome wide.

### SSBP3 is primarily recruited by LDB1

The extraordinarily high co-occupancy of LDB1 and SSBPs suggested that LDB1 might be the primary mode by which SSBPs are recruited to chromatin. We therefore utilized a G1E-ER4 cell line in which endogenous LDB1 was tagged with an mAID-degron^25^. A 4-hour auxin treatment effectively eliminated LDB1 from chromatin as determined by ChIP-seq (Figure 1F). Notably, SSBP3 was also largely eliminated from chromatin following LDB1 depletion (Figure 1F).

G1E-ER4 cells can be induced to undergo erythroid differentiation upon exposure to estradiol^64^. To examine whether the dynamic changes in LDB1 occupancy are accompanied by similar changes in SSBP occupancy, we performed ChIP-seq for LDB1 and all three SSBPs after 12 hours of estradiol treatment. 14,075 and 11,210 LDB1 peaks were gained and lost, respectively (FC ≥ 2, FDR ≤ 0.05) (Figure S1I). Chromatin binding changes of all three SSBPs closely followed those of LDB1 (Figure S1J-M). Together these results suggest that SSBPs are primarily, if not exclusively, recruited to chromatin by LDB1.

### SSBP3 facilitates LDB1 homodimerization to form higher order complexes

Despite the interaction and colocalization of SSBPs with LDB1, how SSBPs affects LDB1 function remained unclear. We began by investigating whether SSBP3 may modulate LDB1 self-association.

First, using the homobifunctional protein crosslinker bis-sulfosuccinimidyl suberate (BS3), shown to crosslink LIM-HD proteins to LDB1 and LDB1 to itself^27^, we investigated whether full-length LDB1 transcribed and translated *in vitro* could homodimerize in solution. Indeed, LDB1, but not SSBP3, formed homodimers (Figure 2A, open arrowhead in lane 3), confirming published results^27^. Homodimerization was inefficient, since at least half of the LDB1 protein remained monomeric under the conditions employed (Figure 2A, lane 3). When SSBP3 was incubated with LDB1 prior to crosslinking, at least two complexes greater than 200kDa were detected by autoradiography (Figure 2B, solid arrowhead in lanes 5-8). Increasing the amount of SSBP3 was associated with a concentration-dependent increase in the abundance of the higher molecular weight complexes with a corresponding decrease in LDB1 monomer.

**Figure 2.**
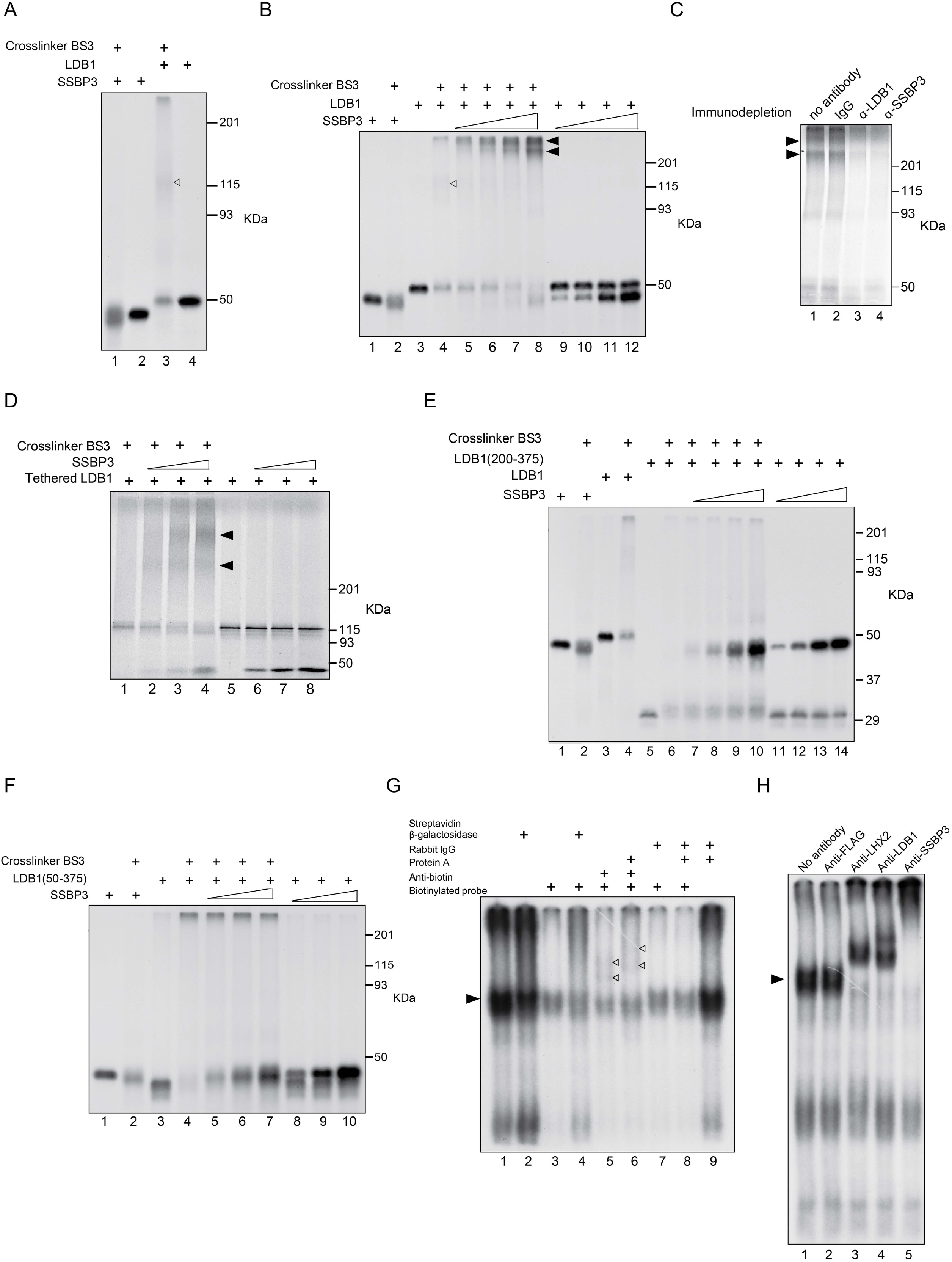
SSBP3 facilitates LDB1 homodimerization. (A) Autoradiograph of SDS-PAGE gel (6%) containing ^35^S-methionine-labeled LDB1 and SSBP3 proteins crosslinked *in vitro* with BS3. Additions to *in vitro* mixture are shown atop each lane. Open arrowhead shows LDB1 homodimer. (B) Autoradiograph of SDS-PAGE gel (8%) containing ^35^S-methionine-labeled constant amount of LDB1 and increasing amount of SSBP3 proteins *in vitro* with BS3 crosslinking. Solid arrowheads show multimers of LDB1-SSBP3. Open arrowhead shows LDB1 homodimer. (C) Autoradiograph of SDS-PAGE-separated proteins from crosslinking assay in which specific antibodies depleted either LDB1 or SSBP3. High molecular weight complexes containing both SSBP3 and LDB1 proteins are shown by solid arrowheads. (D) Autoradiograph of SDS-PAGE (6%) separation of proteins from crosslinking assays using ^35^S-methionine-labeled tethered LDB1 and increasing concentrations of SSBP3. (E) Autoradiograph of ^35^S-methionine-labeled proteins from crosslinking analyses of LDB1 truncation mutant (200-375aa). (F) Autoradiograph of ^35^S-methionine-labeled proteins from crosslinking analyses of LDB1 truncation mutant (50-375aa). (G) Autoradiograph of PAGE-separated protein complexes formed by incubating nuclear extract of αT3-1 cells with ^32^P-labeled DNA probe and biotin-labeled DNA probe. Additions to the incubation mixture are shown atop each lane. The previously reported complex that binds the ^32^P-labeled probe through its LHX2 component is marked by the solid arrowhead. The addition of anti-biotin antibody, either alone or with protein A super-shifted the complex (open arrowheads), indicating the binding of both ^32^P-labeled and biotin-labeled probes. (H) Autoradiograph of EMSA using nuclear extracts from αT3-1 cells and ^32^P-labeled DNA probe. Addition of specific antibodies to components of the LHX2, LDB1, and SSBP3 complex are shown atop each lane. Location of band representing the LHX2, LDB1, and SSBP3 complex bound to ^32^P-labeled probe is shown by solid arrowhead.

To more definitively characterize the components of the complexes formed by BS3 crosslinking, Western blot analysis was carried out using a SSBP3 antibody (Figure S2A) which reacted strongly with un-crosslinked SSBP3 but much less with BS3-crosslinked SSBP3 (Figure S2A). When LDB1 and SSBP3 were crosslinked with BS3, however, SSBP3 was detected in two high molecular weight complexes when higher concentrations of SSBP3 were inputted (Figure S2A, solid arrowheads in lane 7).

Because the LDB1 antibody could not detect crosslinked LDB1 (data not shown), alternate strategies were employed. First, unlabeled methionine was used in place of ^35^S-labeled methionine for LDB1 in *in vitro* translation. When cold LDB1 was then added to ^35^S-labeled SSBP3 in the chemical crosslinking assay, the two high molecular weight LDB1-SSBP3 complexes were detected by autoradiography (Figure S2B, lane 5, solid arrowheads).

To further confirm the identity of these high molecular weight complexes, we, second, carried out immune depletion prior to crosslinking. LDB1 antibody, but not IgG (Figure 2C, lane 2), depleted radiolabeled LDB1 migrating with the ∼ 50kDa marker (lane 3), SSBP3 antibody specifically depleted radiolabeled SSBP3 migrating slightly faster than ∼50kDa (lane 4), and both efficiently depleted the two high molecular weight complexes observed after crosslinking (solid arrowheads in Figure 2C, lanes 1 and 2). Thus, by interacting with homodimeric but not monomeric LDB1, SSBP3 shifts the equilibrium towards LDB1 dimers.

To test whether LDB1 homodimerization is sufficient for SSBP interaction, an expression vector was constructed in which a 22-amino acid glycine linker GT(GGGS)4GGGT was inserted in-frame between two full-length LDB1 peptide-coding sequences. With duplication of LDB1 in a single head-to-tail orientation, intramolecular interactions are greatly favored over intermolecular interactions. When the “tethered” LDB1 dimer (TD-LDB1) was subjected to chemical crosslinking, TD-LDB1 did not form larger complexes (Figure 2D, lanes 1 and 5). However, TD-LDB1 interacted with SSBP3 (Figure 2D, lanes 2-4) such that increasing amounts of SSBP3 led to more TD-LDB1 being incorporated into higher molecular weight complexes (Figure 2D, lanes 2-4 solid arrowheads). TD-LDB1 had a similar turnover rate as native LDB1 (Figure S2C). These results suggest that LDB1 homodimerization is sufficient for SSBP3 binding.

To determine whether homodimerization of LDB1 is required for SSBP interaction, two well characterized dimerization-incompetent LDB1 deletion mutants were employed. The first mutant, LDB1 (200-375) is completely defective in self-association but still capable of binding LIM domain proteins in BS3-mediated crosslinking analysis^27^. We confirmed that LDB1 (200-375) could not form homodimers (Figure 2E, lanes 5 and 6) and showed that it could not bind to SSBP3 (Figure 2E, lanes 7-10). Given that the 199 amino acids deleted in LDB1 (200-375) almost immediately abuts the LCCD, the defect in SSBP interaction could have been due to disruption of this SSBP-LDB1 binding surface. We therefore also tested a second mutant protein with a smaller deletion, LDB1 (50-375) which retains a greater amount of protein sequence N-terminal to the LCCD domain than LDB1(200-375) but is similarly dimerization-defective^65^. Importantly, LDB1 (50-375) was defective in homodimerization and in interaction with SSBP3 (Figure 2F). Thus, LDB1 homodimerization is both necessary and sufficient for stable LDB1-SSBP3 interaction.

These experiments indicate that SSBP3 acts to promote LDB1 homodimerization through its direct contact with LDB1 homodimer, with a resultant shift in equilibrium between monomer and dimer. However, the possibility of SSBP3 contacting LDB1 monomer and causing a conformational change in DD that promotes dimerization cannot be ruled out.

### The SSBP3/LDB1 complex can connect two DNA fragments *in vitro*

To investigate whether SSBP3 may support LDB1-mediated DNA looping *in vitro*, we modified the electrophoretic mobility shift assay (EMSA)^66^ to be able to detect the stable linking of two double-stranded DNA fragments in solution. In this assay, nuclear extract is incubated with a mixture of ^32^P-labeled and biotin-labeled double-stranded DNA probes, and protein-DNA complexes containing both radioactive and biotinylated oligonucleotides are identified from their retardation in non-denaturing polyacrylamide gels with an antibody to biotin. The specificity of the biotin step is proven by a similar shift in the complex’s mobility with addition of streptavidin-β-galactosidase instead.

Using the DNA-binding LIM homeodomain protein LHX2 in place of LMO2, TAL1, and GATA-1 to reduce the complexity of this assay, we demonstrated that incubation of nuclear extract from the SSBP3-, LDB1-, and LHX2-containing αT3-1 pituitary cell line with an empirically determined mixture of biotinylated and radiolabeled probes containing a minimal LHX2 binding element^59^ resulted in a diminution of detectable DNA-linking activity (Figure 2G, solid arrowheads). This decrease is due in part to the fact that the biotinylated probe acts as a cold competitor of radiolabeled probe (Figure 2G, compare lanes 1 and 3). The further retardation of the complexes that were detected in autoradiography, however, by addition of streptavidin-β-galactosidase indicates that they contain both the biotin-labeled and ^32^P-labeled double-stranded probes (Figure 2G, compare lanes 3 and 4).

Use of antibody to biotin in place of streptavidin-β-galactosidase was associated with better detection of super-shifted complexes (Figure 2G, lane 5, open arrowheads), which were further shifted by the addition of protein A (Figure 2G, compare arrowheads in lanes 5 and 6). Lanes 7, 8 and 9 in Figure 2G constitute controls showing the absence of the shifted complexes when rabbit IgG was used in place of biotin antibody. Finally, antibody super-shift analysis indicates that this DNA-linking complex contains LHX2, LDB1, and from the different mammalian SSBPs possible, predominantly SSBP3 (Figure 2H).

From these results, we suggest that an SSBP3-, LDB1-, and LHX2-containing complex can link two DNA elements in solution, simulating the looping of two DNA elements *in vivo*. These findings begin to delineate a mechanism by which SSBP3 promotes LDB1 mediated DNA looping *in vivo*.

### *Ssbp3* but not *Ssbp2* and *Ssbp4* is essential for erythroid cell viability

To examine to what extent SSBP2/3/4 have distinct or overlapping functions, we attempted to generate knockout cell lines for all three via Cas12a-based genome editing^67^. Specifically, we used three guide RNAs simultaneously targeting the conserved N-terminal LUFS domains of SSBP2/3/4 (Figure S3A). Among 29 clones examined, the only ones we were able to obtain contained mutations in *Ssbp2* and *Ssbp4*, while leaving *Ssbp3* intact (Figure S3B). Clone #1 harbors short in-frame deletions in both *Ssbp2* alleles. Clone #2 contains a frameshift mutation in one allele and an in-frame deletion in the other. These in frame deletions reside within the first alpha helix of the LUFS domain, likely impacting SSBP2 protein stability and functionality^68^ (Figure S3B). Both clone #1 and #2 harbor homozygous frameshift mutations in *Ssbp4*. Clone #3 is a compound knockout of *Ssbp2/4* as it contains frameshift mutations in both *Ssbp2* (19 bp deletion) and *Ssbp4* (23 bp deletion) (Figure S3B).

We then used CRISPR/Cas9 in an attempt to knock out *Ssbp3* in both *Ssbp2/4* knockout clone #3 and the parental G1E-ER4 cells. Cas9 targeting the SSBP3 LUFS domain caused cell death both in clone #3 and the parental G1E-ER4 cells. We isolated the surviving cells, purified genomic DNA and performed Sanger sequencing on the target region and found they contained wild type *Ssbp3* alleles. Similar results were obtained with a second guide RNA targeting the C-terminal single stranded DNA binding domain of SSBP3. These results suggest that *Ssbp3* is essential for viability of G1E-ER4 cells.

### LDB1 abundance and chromatin binding are robust to combined loss of all SSBPs

SSBP proteins were suggested to regulate the abundance of LDB1 by shielding it from degradation^55,56^. Since SSBP2/4 combined loss was tolerated by G1E-ER4 cells, we examined whether it affected LDB1 abundance. In clones #1, #2, and #3, the total SSBP2/4 protein levels were reduced as determined by Western blot (Figure 3A) with an antibody that reacts with both SSBP2/4 (Figure S3C, D, E). We did observe residual bands of unknown identity with similar molecular weights that do not represent SSBP2/3/4 because SSBP2/4 were knocked out and because the antibody does not react with SSBP3. Importantly, total protein levels of LDB1 and LMO2 were unchanged (Figure 3A). To examine whether SSBP2/4 may affect protein levels on chromatin, we performed ChIP-seq for LDB1, LMO2, and SSBP3 in SSBP2/4 depleted cells (clone #3) and found essentially no changes when compared to controls (Figure 3B). At LDB1 bound sites H3K27ac, a mark of active chromatin, also remained unchanged in SSBP2/4 depleted cells, indicating that any enhancer activity at LDB1 occupied enhancers was unperturbed (Figure 3B). Lastly, RNA-seq revealed that SSBP2/4 depletion had minimal effect on gene expression in undifferentiated condition (minus estradiol) (Figure S3F and G). Minor changes in gene expression were observed in differentiated condition (plus estradiol) (Figure S3F and G). Whether the minor changes in gene expression were due to direct or secondary effects remains unknown. In sum, SSBP2 and SSBP4 are dispensable for LDB1 protein abundance, chromatin binding, enhancer activity, and largely also for gene transcription, leading us to focus on SSBP3 as the primary functional SSBP in erythroid cells.

**Figure 3.**
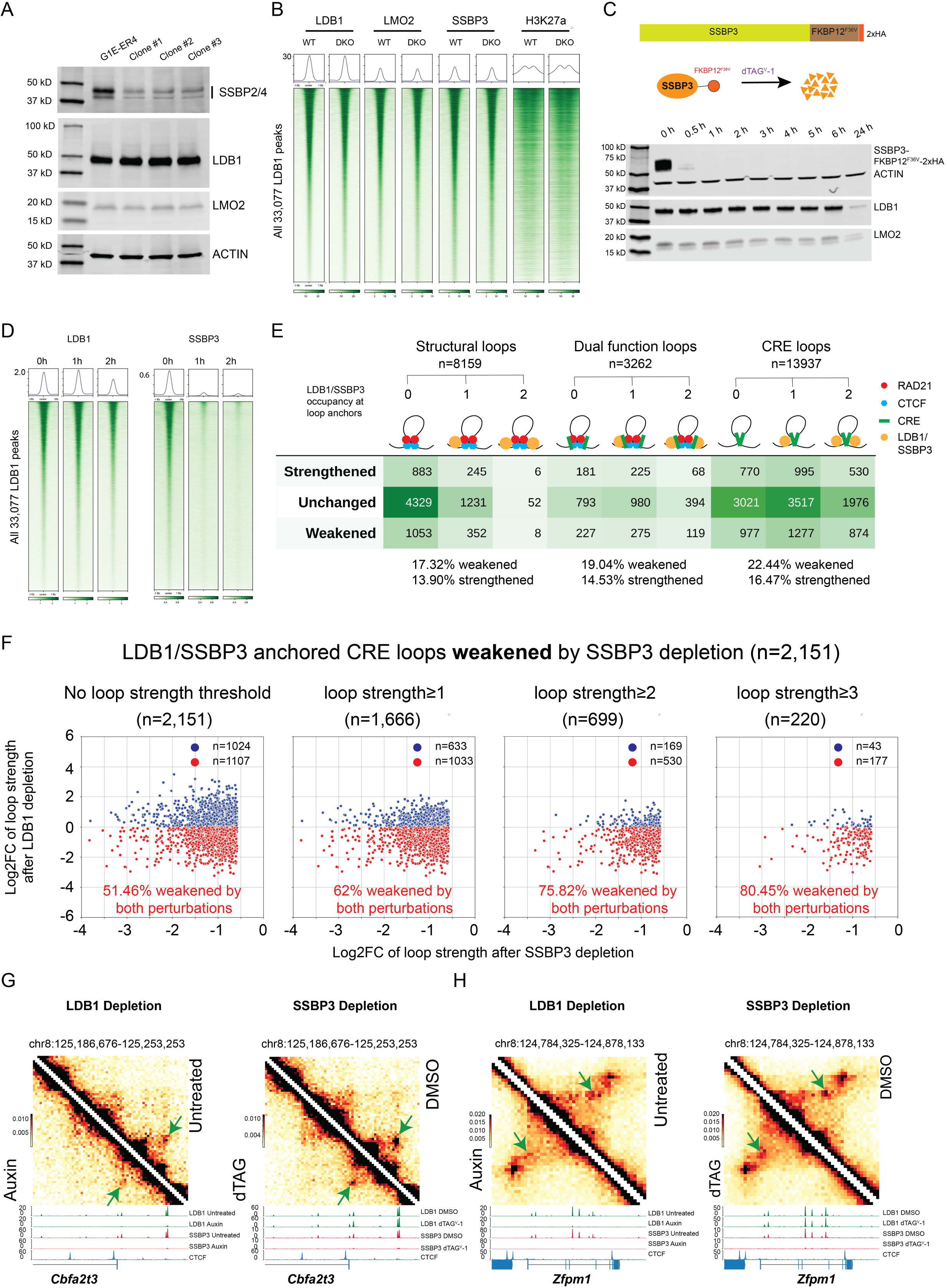
Acute depletion of SSBP3 in *Ssbp2/4* knockout cells affects chromatin looping without altering LDB1 protein abundance and chromatin binding. (A) Western blot with indicated antibodies in G1E-ER4 and *Ssbp2/4* hypomorphic (clone #1 and #2) and knockout (clone #3) cells. ACTIN was used as a loading control. Western blotting was performed at least three times and representative result is shown. (B) Heatmaps centered on all LDB1 peaks showing LDB1 (n=4), LMO2 (n=3), SSBP3 (n=2), and H3K27ac (n=2) were unchanged in *Ssbp2/4* knockout cells compared to G1E-ER4 cells. (C) Upper: diagram showing FKBP12^F36V^-2xHA was knocked in at the C terminus of SSBP3; Middle: diagram showing dTAG^V^-1 induced degradation of SSBP3-FKBP12^F36V^-2xHA; Lower: Western blot analysis of total proteins from SSBP3-FKBP12^F36V^-2xHA cells treated with 500 nM dTAG^V^-1 over a time course. HA antibody was used to confirm the degradation of SSBP3-FKBP12^F36V^-2xHA protein. LDB1 and LMO2 antibodies were used to show their protein abundance were unaffected within 6h but were reduced at 24h. ACTIN was used as a loading control for the inputs. WB was performed at least three times and representative result is shown. (D) Heatmaps showing genome wide LDB1 (n=3) binding was unaffected after 1h SSBP3 (n=3) depletion and was mildly decreased after 2h SSBP3 (n=3) depletion. (E) Upper: Diagram showing three categories of loops: structural loops, dual function loops, and CRE loops, which were further stratified by LDB1/SSBP3 occupancy at loop anchors. Lower: Differential analysis for the indicated loop types after SSBP3 depletion. (F) Scatterplots showing loops that were weakened by both SSBP3 depletion and LDB1 depletion (red dots) and loops that were weakened by SSBP3 depletion but not by LDB1 depletion (blue dots). The more stringent threshold of loop strength (observed over locally adjusted expected value) is used, the higher overlapping rate is observed (G-H) Pairwise comparison between Micro-C contact heatmaps from LDB1 depletion and SSBP3 depletion (n=2) at *Cbfa2t3* (G) and *Zfpm1* (H) loci, showing LDB1/SSBP3 dually anchored CREs loops (marked by green arrows) were weakened by both depletions. Untreated and 4h auxin treated LDB1 (n=2) and SSBP3 (n=2) ChIP-seq tracks were shown below the LDB1 Micro-C heatmaps. 1h DMSO treated and 1h dTAG^V^-1 treated LDB1 (n=3) and SSBP3 (n=3) ChIP-seq tracks were added below the SSBP3 Micro-C heatmaps. CTCF (n=2) ChIP-seq track in G1E-ER4 were added to mark nearby structural loops.

To test whether SSBP3 regulates LDB1 abundance, we knocked-in a FKBP12^F36V^-2xHA tag^69^ at the C-terminus of SSBP3 in the SSBP2/4 depleted cells (clone #3) (Figure 3C). A time series of dTAG^V^-1 treatments showed that SSBP3 could be efficiently depleted as early as 0.5 hours (Figure 3C). Global LDB1 and LMO2 protein levels remained constant for up to 6 hours, suggesting that within this period, SSBP3 was dispensable for LDB1 stability (Figure 3C). However, at 24 hours, LMO2 and LDB1 levels were diminished. Whether this was due to protein turnover or secondary effects is unknown. To examine whether chromatin-bound SSBP3 was depleted after dTAG^V^-1 treatment, and whether LDB1 binding was affected by SSBP3 depletion, we performed ChIP-seq for SSBP3 and LDB1 at 0h, 1h, and 2h of dTAG^V^-1 treatment. We found that while SSBP3 was depleted from chromatin after 1h dTAG^V^-1 treatment (Figure 3D), LDB1 binding was stable after 1h of dTAG^V^-1 treatment (107% of 0h, based on aggregate ChIP-seq signals) and showed only a slight reduction after 2 hours (73% of 0h) (Figure 3D). Based on these observations, SSBP3 does not appear to regulate LDB1 protein abundance, at least within a short time frame.

### SSBP3 is required for enhancer-promoter (E-P) looping

LDB1 contributes widely to regulatory connectivity among enhancers and promoters. Having established a time point at which we could deplete SSBP3 while leaving LDB1 chromatin binding intact enabled us to assess a direct role of SSBP3 in LDB1-mediated chromatin looping.

We performed Micro-C^70^ using 1-hour DMSO (control) or dTAG^V^-1 treated SSBP3-dTAG cells (in the SSBP2/4 knockout clone #3 background). Two biological replicates generated a total of 1.431 billion and 1.402 billion valid cis contacts for the DMSO and dTAG^V^-1 treatment conditions, respectively (Supplementary Table 1). Acute depletion of SSBP3 did not result in changes in genome-wide contact decay curves (Figure S4A), A/B compartmentalization (Figure S4B), and TAD boundary insulation (Figure S4C).

To study the effects of SSBP3 loss on chromatin looping we employed the loop callers Cooltools^71^ and Mustache^72^. Mustache called more loops at higher resolution (1kb, and 2kb; here, resolution refers to the bin size used to divide the genome when building the Micro-C contact matrix), while Cooltools called more loops at lower resolution (10kb) (Figure S4D). Since Mustache was able to call smaller loops (closer to the diagonal in the Micro-C contact heatmap) that were not called by Cooltools (two representative regions are shown in Figure S4E), and since E-P and E-E loops tend to be shorter than, for example, CTCF/cohesin- mediated structural loops, we used the loop calls from Mustache.

We identified 9,385 loops at 1kb resolution, 16,950 loops at 2kb resolution, 15,632 loops at 5kb resolution, and 8,548 loops at 10kb resolution by merging loops called separately under DMSO and dTAG^V^-1 treatment conditions (FDR ≤ 0.05). In total, we identified 40,194 loops by merging loop calls from all resolutions and both treatment conditions (Note that the total loop number is not the sum of the numbers from each resolution. When loop anchors of two loops overlap, only the loop in finer resolution is counted). Loop strength was quantified by measuring the observed contact frequency within the peak pixel, divided by a locally adjusted expected value, under both DMSO and dTAG^V^-1 conditions. Changes in loop strength were thresholded at a fold change of ≥1.5, ≤0.67 (log2FC cutoff of ±0.585) and were assessed at each resolution at which the loops were called. After 1 hour of dTAG^V^-1 treatment, we observed more weakened than strengthened loops at all resolutions (Figure S4F). In total, we identified 8,343 weakened loops, 6,436 strengthened loops, and 25,415 unchanged loops (Figure S4G).

We stratified loops affected by SSBP3 depletion into 1) structural loops - loops co-occupied by CTCF/RAD21 at both anchors, with H3K27ac present at one or neither anchor, 2) dual function loops - loops where CTCF/RAD21 were occupied at both anchors, with H3K27ac present at both anchors, and 3) Cis-Regulatory Element (CRE) loops - loops with H3K27ac present at both anchors but excluding loops with CTCF/RAD21 at both anchors (Figure 3E). For these classifications we used 10kb windows centered on the central pixels of the loop anchor. Loop stratification was performed without loop strength thresholding.

Based on this stratification, 65.8% of CRE loops (9,169 out of 13,937), 63.2% of dual function loops (2,061 out of 3,262), and only 23.2% of structural loops (1,894 out of 8,159) had LDB1/SSBP3 co-bound at one or both loop anchors (Figure 3E). These results suggest that the majority of CRE loops are bound by LDB1/SSBP3 at one or both anchors.

22.44% of CRE loops (3,128 out of 13,937) were weakened, and 16.47% of CRE loops (2,295 out of 13,937) were strengthened after SSBP3 depletion. The majority of CRE loops (61.09%, 8,514 out of 13,937) were unaffected (Figure 3E). A large fraction of loops with SSBP3 at their anchors were insensitive to SSBP3 depletion. Residual SSBP3 binding after 1h depletion may still be capable of maintaining the loops. It is also possible that the large window size (10kb) assigned to loop anchors allows for SSBP3 occupancy being called without a direct involvement at these anchors. In addition, SSBP3 at these sites may control loops that were too short to be resolved by Micro-C. A case in point, when examining LDB1 dependent loops, use of higher resolution techniques increased the number of called loops and increased the correlation between occupancy and looping function^25^.

LDB1/SSBP3 anchored CRE loops accounted for 68.77% of total weakened CRE loops (2,151 out of 3,128) and 66.45% of total strengthened CRE loops (1,525 out of 2,295) (Figure 3E), suggesting SSBP3 depletion may directly affect these loops. However, we also found 31.23% of weakened (977 out of 3,128) and 33.55% of strengthened (770 out of 2,467) CRE loops that lacked measurable LDB1/SSBP3 occupancy at their anchors.

Using H3K27ac/H3K4me1 for CRE annotation produced similar loop stratification results as using H3K27ac alone (Figure S4H and I).

To determine the type of CRE interactions dependent on LDB1/SSBP3, we further subdivided CRE loop anchors into promoters (P; 2kb region upstream of TSS) and enhancers (E; CREs not overlapping with promoters). The majority of LDB1/SSBP3 co-dependent CRE loops were E-E loops with LDB1/SSBP3 bound at both anchors (Figure S4J), very similar to LDB1-dependent loops^25^.

19.04% of dual function loops (621 out of 3,262) were weakened and 14.53% of dual function loops (474 out of 3,262) were strengthened after SSBP3 depletion. LDB1/SSBP3 anchored dual function loops accounted for 63.45% of total weakened dual function loops (394 out of 621) and 61.81% of total strengthened dual function loops (293 out of 474) (Figure 3E). 36.55% of weakened (227 out of 621) and 38.19% of strengthened (181 out of 474) dual function loops lack LDB1/SSBP3 occupancy.

Analysis of structural loops (as defined as CTCF/cohesin flanked loops) revealed that SSBP3 depletion led to a weakening of 17.32% (1,413 out of 8,159) and a strengthening of 13.90% (1,134 out of 8,159). A substantial proportion of structural loops (68.78%; 5,612 out of 8,159) were not affected by SSBP3 depletion. Since our recent study showed the genome wide binding of CTCF and RAD21 remained stable upon LDB1 depletion^25^, it is unlikely that SSBP3 loss alters CTCF or RAD21 occupancy. Instead SSBP3 loss may influence structural loops via the effects on adjacent loops (Figure 3E).

Based on current sequencing depth (around 1.4 billion valid contacts per condition), a portion of the differential loops identified are likely due to experimental variation. However, we observed a high concordance among each replicate (Figure S5A-D).

Differential loop calls using a range of log2FC cutoffs consistently showed that CRE loops are most vulnerable to SSBP3 depletion (Figure S5E).

Given the high degree of overlap in chromatin occupancy between LDB1 and SSPB3, we asked if their loss affects similar cohorts of loops. First, we assessed whether the 2,151 LDB1/SSBP3 anchored CRE loops that were weakened upon SSBP3 depletion were also weakened upon LDB1 depletion.

To further address potentially confounding effects of experimental variation, we applied thresholds for baseline loop strengths which lowered the number of called loops but increased the concordance between the effects of LDB1 and SSPB3 depletion. Specifically, we quantified loop strength by measuring the observed contact frequency within the peak pixel divided by locally adjusted expected values. We found that, depending on loop strength, up to 80.45% of LDB1/SSBP3 anchored loops that require SSBP3 were weakened upon LDB1 depletion (Figure 3F). Conversely, we found that up to 76.11% of loops that are lost upon LDB1 depletion were also weakened by SSBP3 depletion (Figure S7). In both cases, we observed that strong CRE loops tended to have high LDB1/SSBP3 binding intensities at their loop anchors, which may explain why they were more sensitive to both perturbations as compared to weak CRE loops (Figure S6 and 7).

We also determined whether CRE loops that were strengthened upon SSBP3 loss were also strengthened upon LDB1 depletion. Up to 83.2% of loops were strengthened by both SSBP3 depletion and LDB1 depletion (Figure S8).

Together, our results indicate that LDB1 and SSBP3 are involved in the maintenance of a highly overlapping set of chromatin loops.

Approximately 20% of loops that were weakened by SSBP3 depletion were not weakened by LDB1 depletion, even when selecting the strongest loops in both datasets (Figure 3F). It is possible that the residual LDB1 and SSBP3 binding after 4h auxin treatment (to deplete LDB1) may maintain these loops. Indeed, when we examined residual LDB1 and SSBP3 ChIP-seq signals after 4h auxin treatment at these loop anchors, we found they had higher residual LDB1 and SSBP3 binding after 4h auxin treatment (Figure S6). Conversely, loops that were weakened by LDB1 depletion but not by SSBP3 depletion had higher LDB1 and residual SSBP3 binding at loop anchors after 1h dTAG^V^-1 treatment (Figure S7).

Representative examples of loop changes are shown at two loci: *Cbfa2t3* (Figure 3G) and *Zfpm1* (Figure 3H). We observed under both depletion conditions, two LDB1/SSBP3 anchored loops were similarly weakened (Figure 3G and H). Additional examples that were weakened by both LDB1 depletion and SSBP3 depletion are shown in Figure S9.

In sum, SSBP3 and LDB1 function in a highly coordinated and mutually dependent manner during chromatin loop formation.

### SSBP3 and LDB1 regulate overlapping sets of genes

SSBP3 and LDB1 may regulate gene expression predominantly via chromatin looping or via looping-independent mechanisms. The former scenario would suggest very similar gene expression changes upon depletion whereas the latter might reveal transcription factor dependent differences, for example, via factor-specific co-factors^43,73–76^. We performed precision nuclear run-on sequencing (PRO-seq) upon 1 hour of SSBP3 depletion^77^. In total, we identified 566 downregulated genes and 173 upregulated genes after SSBP3 depletion (FC ≥ 1.5, FDR ≤ 0.05) (Figure 4A). Several gene expression changes were validated by primary transcript RT-qPCR (Figure S10A). We intersected our PRO-seq data with previously acquired TT-seq data generated upon LDB1 depletion^25^ (Figure 4B and C). We found 83.83% of the downregulated genes following SSBP3 depletion were also downregulated following LDB1 depletion; 92.85% of the upregulated genes following SSBP3 depletion were also upregulated following LDB1 depletion (Figure 4B). In the converse analysis, 94.58% of the downregulated genes following LDB1 depletion were also downregulated following SSBP3 depletion; 75.54 % of the upregulated genes following LDB1 depletion were also upregulated following SSBP3 depletion (Figure 4C). Two examples of downregulated genes and their SSBP3 and LDB1 occupancy profiles are shown in Figure 4D and E. Hence, despite the different methods used to quantify gene expression and the different depletion durations, SSBP3 and LDB1 regulate a highly overlapping group of genes, suggesting that these proteins function by the same mechanism, in this case architecturally, and not via factor-selective co-factor interactions. These findings also indicate that LDB1 is incapable of maintaining gene expression in the absence of SSBP3 in spite of its maintained chromatin occupancy.

**Figure 4.**
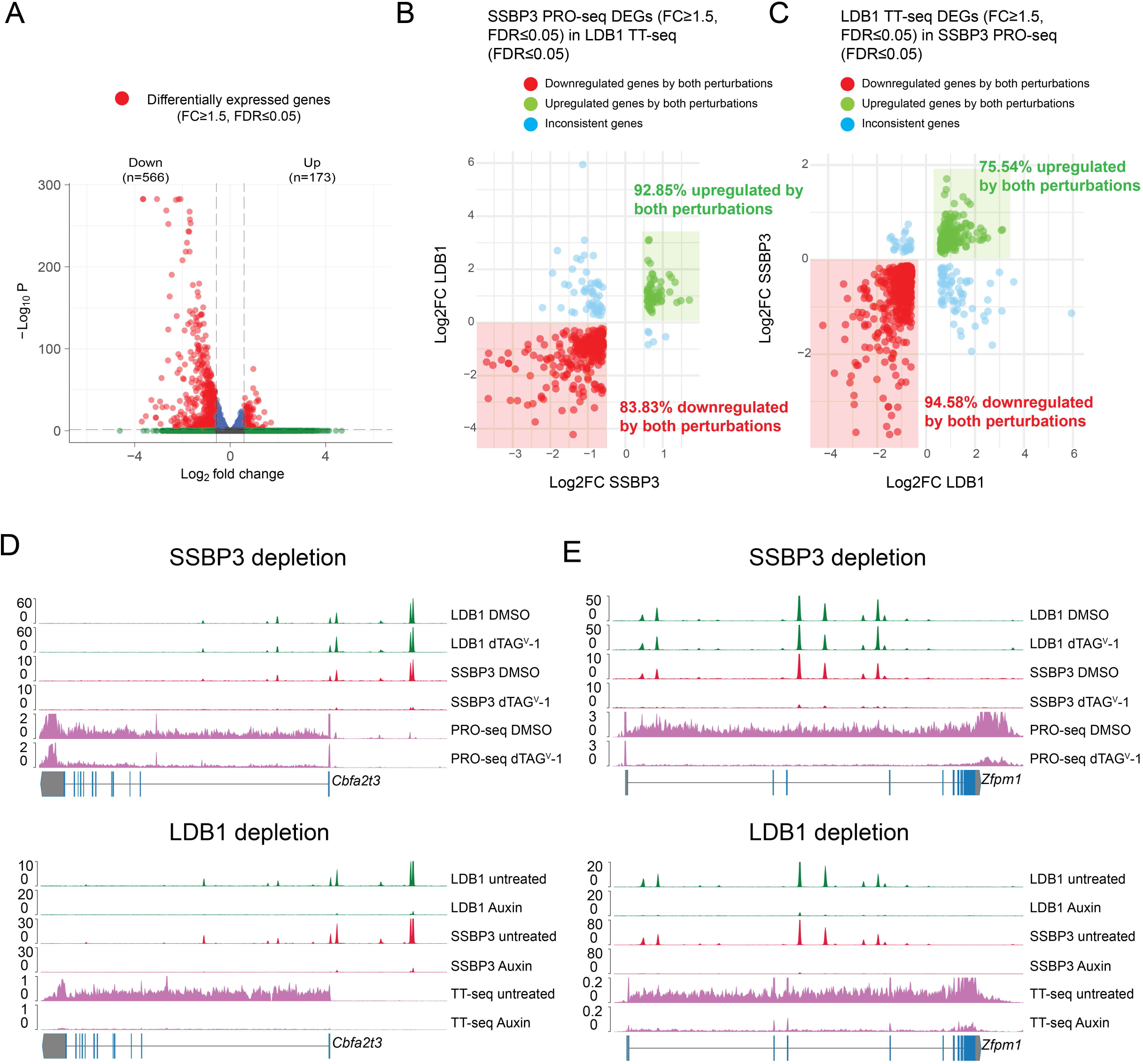
Acute depletion of SSBP3 disrupts nascent transcription of hundreds of genes, mirroring the effects of LDB1 depletion. (A) Volcano plot showing differentially expressed genes between 1h DMSO (n=3) and dTAG^V^-1 (n=3) treatment as revealed by PRO-seq. (B) Scatter plot showing a large fraction of the down- or up-regulated genes upon SSBP3 depletion (PRO-seq) were also down-or up-regulated by LDB1 depletion (TT-seq). (C) Scatter plot showing a large fraction of the down- or up-regulated genes upon LDB1 depletion (TT-seq) were also down- or up-regulated by SSBP3 depletion (PRO-seq) (D-E) Pairwise comparisons of nascent transcription for *Cbfa2t3* (D) and *Zfpm1* (E) following SSBP3 or LDB1 depletion. For SSBP3 depletion, PRO-seq (n=3) along with LDB1 (n=3) and SSBP3 (n=3) ChIP-seq after 1h DMSO or dTAG^V^-1 treatment was shown; For LDB1 depletion, TT-seq along with LDB1 (n=2) and SSBP3 (n=2) ChIP-seq before and after 4h auxin treatment was shown.

To examine the relationship between changes in chromatin looping and transcription, we intersected LDB1/SSBP3-anchored lost (thresholded at Log_2_FC ≤ −0.585) or gained (thresholded Log_2_FC ≥ 0.585) CRE loops with a 2 kb region centered on the transcription start site (TSS) of differentially expressed genes with an FDR ≤ 0.05. We observed that the greater the number of weakened loops linked to a TSS, the greater the downregulation of transcription tended to be (Figure S10B). Using all genes instead of just differentially expressed genes gave similar correlations (Figure S10B).

As previously observed^25^, changes in loop strength and transcription were not correlated (Figure S10C), possibly reflecting the nonlinear relationship between enhancer–promoter contacts and transcriptional output^78^. Nevertheless, up to 44.78% of LDB1/SSBP3 anchored CRE loops weakened upon SSBP3 depletion contacted a downregulated gene (FDR ≤ 0.05) (Figure S10C).

In summary, our results demonstrate that SSBP3 and LDB1 not only colocalize throughout the genome and regulate similar groups of CRE loops, but also influence transcription of similar sets of genes. Notably, the association of downregulated genes with weakened CRE looping further supports the notion that SSBP3, in complex with LDB1, regulates gene expression through its architectural role.

### E-P looping and transcription activation via targeted tethering of SSBP3

We previously engineered a chromatin loop at the endogenous β-globin locus in G1E cells^20^. G1E cells are erythroblasts in which the erythroid transcription factor GATA1, which normally recruits LDB1 to the β-globin promoter, has been knocked out and thus fails to form a chromatin loop between the LCR and the β-globin promoter. Expression of an artificial zinc finger (ZF) designed to bind the β-globin promoter fused to LDB1 bypassed the requirement of GATA1 and restored this loop, supporting a direct role of LDB1 in loop formation^20^. If SSBP3 is an essential component of the LDB1 protein complex, tethering of SSBP3 to the β-globin promoter is expected to similarly form this loop (Figure 5A). We fused the same ZF to full length HA-tagged SSBP3 (SSBP3-FL) or SSBP3 N terminal LUFS domain (SSBP3-N, 1-98 aa), and introduced these constructs into G1E cells (Figure S11A). Anti-HA ChIP-qPCR confirmed that ZF-SSBP3-FL and ZF-SSBP3-N bound the β*-major* promoter (Figure 5B). ZF-SSBP3-FL and ZF-SSBP3-N increased primary β*-major* transcripts 243- and 277- fold, respectively, compared to control G1E cells, indicating strong enhancer-dependent and hence loop-initiated transcription^20^ (Figure 5C). 4C experiments demonstrated that ZF-SSBP3-FL and ZF-SSBP3-N induced interactions between the *β-major* promoter and the LCR (Figure 5D).

**Figure 5.**
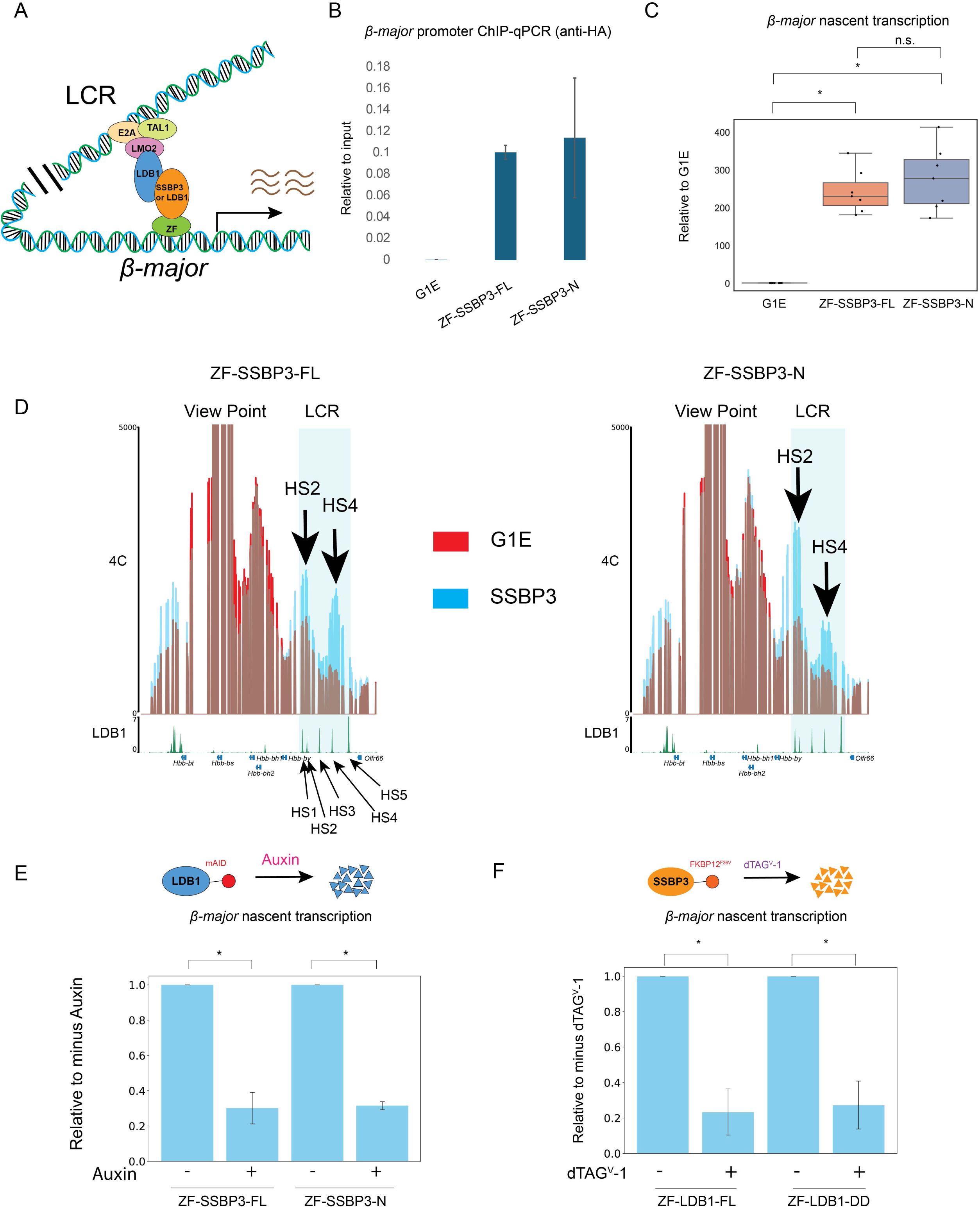
Forced-tethering SSBP3 to the β-Globin promoter triggers spatial proximity with the LCR. (A) Diagram showing the strategy of tethering SSBP3 to the β*-major* promoter using an artificial zinc finger protein to mediate E-P looping and transcriptional activation. (B) HA ChIP-qPCR (n=2) showing ZF-SSBP3-FL and ZF-SSBP3-N binds β*-major* promoter. (C) β*-major* nascent transcripts measured by RT-qPCR (n=7) in G1E cells and derivatives expressing ZF-SSBP3-FL and ZF-SSBP3-N. The box represents the interquartile range (IQR), with the line inside indicating the median. Whiskers extend to the smallest and largest values within 1.5 times the IQR. Statistical significance was assessed using the two-sided Mann-Whitney U test. * *p*≤0.001. (D) 4C-seq in G1E cells expressing ZF-SSBP3-FL (n=2) and ZF-SSBP3-N (n=2). The β*-major* promoter served as the viewpoint. The LCR enhancer is highlighted in cyan. LDB1 ChIP-seq track in undifferentiated G1E-ER4 was added below each 4C track. (E) Upper: diagram showing the strategy to degrade LDB1. Lower: β*-major* nascent transcripts levels as measured by RT-qPCR (n=4) in LDB1-AID cells expressing ZF-SSBP3-FL or ZF-SSBP3-N with or without 4h auxin treatment. The box represents the interquartile range (IQR), with the line inside indicating the median. Whiskers extend to the smallest and largest values within 1.5 times the IQR. Statistical significance was assessed using the two-sided Student T test. * *p*≤0.001. (F) Upper: diagram showing the strategy to degrade SSBP3. Lower: β*-major* nascent transcripts levels as measured by RT-qPCR (n=6 for ZF-LDB1-FL, n=5 for ZF-LDB1-DD) in SSBP3-dTAG cells expressing ZF-LDB1-FL or ZF-LDB1-DD with or without 1h dTAG^V^-1 treatment. The box represents the interquartile range (IQR), with the line inside indicating the median. Whiskers extend to the smallest and largest values within 1.5 times the IQR. Statistical significance was assessed using the two-sided Student T test. * *p*≤0.001.

We observed that HS2 and HS4 exhibited slightly different contact increases in response to the introduction of ZF-SSBP3-FL or ZF-SSBP3-N (Figure 5D). The reason for this difference is unknown.

To examine whether ZF-SSBP3 functions through LDB1, we introduced ZF-SSBP3-FL and ZF-SSBP3-N into LDB1-AID cells^25^ (Figure 5E). After 4-hour auxin treatment, to deplete LDB1, the induction of *β-major* nascent transcription was drastically reduced (Figure 5E).

To test whether, conversely, LDB1-forced E-P looping requires SSBP3, we expressed ZF-LDB1-FL and ZF-LDB1-DD (DD refers to the LDB1 N terminal dimerization domain) into SSBP3-dTAG cells. After one-hour dTAG^V^-1 treatment, *β-major* nascent transcript levels were strongly diminished (Figure 5F). Additionally, 4C experiments revealed weakened interactions between the *β-major* promoter and the LCR (Figure S11B). Thus, the use of a defined engineered system demonstrates that SSBP3 and LDB1 depend on each other to form looped contacts at the β-globin locus.

## Discussion

We report that SSBPs exert critical functions in chromosomal folding as part of the LDB1 architectural complex. 1. SSBP2/3/4 are highly co-localized with LDB1 genome wide. 2. LDB1 depletion leads to an almost complete loss of SSBP2/3/4 chromatin occupancy, suggesting that LDB1 is the predominant, if not only, tether of SSBPs to chromatin, and that single stranded DNA has little if any influence. 3. Acute depletion of SSBP3 on a SSBP2/4 deficient background and acute depletion of LDB1 affect highly overlapping sets of chromatin loops and genes. 4. SSBP3 stabilizes LDB1 dimers *in vitro*, providing a possible mechanism by which it contributes LDB1 loop formation. 5. Engineered chromatin loops revealed a mutual dependence of SSBP3 and LDB1 in establishing long range chromatin contacts.

In erythroblasts, all three members of the SSBP family are expressed but only SSBP3 is essential for cell viability. SSBP3 can therefore compensate for the loss of the other family members but not vice versa. The physical interaction between SSBPs and LDB1 were observed, and experiments based on prolonged perturbations suggested that SSBPs predominantly function by preventing LDB1 degradation^55,56,58–60^. However, it is a common observation that removal of a component from a molecular assembly can destabilize the entire complex, leaving open the possibility that SSBPs mediate LDB1 function in additional ways. To address this question, we employed an acute degradation system to remove SSBP3 in a manner leaving LDB1 protein levels and chromatin occupancy intact. This revealed a widespread role for SSBP3 in supporting LDB1’s architectural function. SSBP3 very likely acts directly on LDB1 as its role was revealed upon depletion for less than one hour.

Even though the SSBPs earned their name because of their ability to bind single stranded DNA *in vitro*^33^, there is little evidence to support such a function *in vivo*. Loss of LDB1 globally abrogates SSBP3 chromatin occupancy but is not expected to affect the occurrence of single stranded DNA. Therefore, single stranded DNA is a minor if any contributor to SSBP recruitment to chromatin *in vivo*.

Loss of SSBP3 and LDB1 affect highly overlapping sets of loops. Gene expression changes incurred by the absence of SSBP3 matched those observed upon LDB1 loss even though LDB1 chromatin occupancy was maintained. Moreover, forced chromatin contacts instigated by tethering of LDB1 or SSBP3 to the β-globin gene revealed a mutual dependence for loop formation in a highly defined and reductionist system. Collectively, these findings indicate that LDB1 and SSBP3 function in a highly coordinated manner as part of multi-component assemblies.

We previously reported that the majority of LDB1-dependent contacts form independently of CTCF and cohesin^25^. First, LDB1 and CTCF/cohesin rarely co-localize. Second, acute depletion of LDB1 has little impact on CTCF/cohesin chromatin occupancy, and vice versa. Third, cohesin-driven chromatin loop extrusion is not required for the majority of LDB1 mediated contacts^25^. The close functional and physical relationship between LDB1 and SSBP3 suggests that SSBP3 is likewise a CTCF/cohesin-independent architectural factor with the sole function to support LDB1. Notably, enhancer activity as reflected by H3K27ac is maintained in acutely LDB1-depleted cells. This indicates that LDB1 and SSBP3 regulate gene expression predominantly via their ability to spatially connect regulatory elements rather than acting as conventional transcription factors.

An open question is why a subset of loops was gained upon SSBP3 loss. We speculate that enhancers that are no longer connected by the LDB1/SSBP3 complex engage in *de novo* contacts via alternate mechanisms that are normally disfavored. In support of this possibility, the gained loops involved enhancer-enhancer contacts. LDB1/SSBP3 complexes might thus constrain “illegitimate” chromatin contacts.

Why a fraction of loops with LDB1/SSBP3 at their anchors remained intact following SSBP3 loss is unclear. Residual SSBP3 binding after 1h depletion may sustain these loops. Lastly, there were loops that were reduced or lost upon SSBP3 degradation even though they ostensibly lacked LDB1/SSBP3 at their anchors.

This could be due to false negative ChIP results. As shown in Fig.S1, changes in experimental conditions vastly increased the number of SSBP3 ChIP peaks. It is possible that further optimization of the ChIP protocol would result in even greater numbers of peaks.

At the *Car2* locus, LDB1 was reported to form a loop by directly binding via its LCCD domain to CTCF, and deletion of the LCCD domain diminished LDB1’s ability to form this loop, implicating CTCF as a partner at heterotypic loop anchors^31^. However, since deletion of the LCCD domain might also affect SSBP3 binding, it remains possible that loop loss might be instead caused by the possible reduced ability to bind SSBP3. Consistent with this possibility, our PRO-seq data showed that *Car2* gene expression was strongly downregulated within one hour of SSBP3 depletion (Log2FC = −2.08; FDR ≈ 0). Moreover, most LDB1-CTCF anchored loops were resistant to CTCF depletion (Nicholas Aboreden, unpublished observation), suggesting that CTCF even though present at some loop anchors may not be the driving force for such contacts.

Finally, our studies provide a possible mechanism by which SSBP3 promotes the chromatin looping function established for LDB1. Specifically, *in vitro* studies showed that SSBP3 binds preferentially to LDB1 dimers, shifting the equilibrium from monomeric to dimeric forms of LDB1. This in turn would facilitate contacts between LDB1 bound loop anchors. Of note, recent experiments suggest that protein assemblies containing LDB1 and associated proteins can be exceedingly stable and are linked to enhancer hubs, at least when studied in non-dividing olfactory neurons^79^. Whether similarly stable structures form in cells with different milieus and that are rapidly dividing will await live cell imaging studies.

LDB1 may not only form dimers but also trimers, while LDB2 (a homolog of LDB1) can even form octamers^29^. Whether SSBP3 can also stabilize higher order LDB1 multimers remains to be tested.

In contrast to published results showing that SSBPs can dimerize^52–54,68^, we did not observe SSBP3 dimerization in solution. However, one difference between our and others’ results is that we used full length SSBP3, while others only used only the N-terminal LUFS domain of SSBP protein^52–54,68^.

In sum, SSBP3 and likely other SSBP proteins are essential components of architectural LDB1 complexes which establish spatial connections between enhancers and promoters to drive gene expression. Our work also raises the possibility that SSBP proteins diversify LDB1 functions depending on the exact SSBP partner that LDB1 interacts with in different cellular backgrounds.

### Limitations of the study

At the resolution afforded by Micro-C with the number of valid pairs, we failed to detect a short range loops, which might affect the correlation between loop calls and LDB1/SSBP3 occupancy.

*In vitro* protein binding studies may have missed higher order LDB1/SSBP3 assemblies due to either lack of additional required components or the detection limits of the gel electrophoresis used.

## Methods

### Cell culture

G1E-ER4 is derived from the murine erythroblast cell line G1E^57^. All genetically modified cell lines used in this study are derivates of G1E-ER4. Cell lines were grown in IMDM + 15% FBS, penicillin/streptomycin, Kit ligand, monothioglycerol and erythropoietin in a standard tissue culture incubator at 37 °C with 5% CO_2_. Cells grew at a density of ∼1 million per ml. To differentiate cells, they were seeded at a density around 0.5 million per ml and 100 nM estradiol was added to medium for 12h or 24h. For dTAG based acute depletion assay, cells were seeded at a density of ∼1 million per ml and 500 nM dTAG^V^-1 was added for the indicated durations.

HEK293T (ATCC) and αT3-1 cells were cultured in Dulbecco’s modified Eagle’s medium containing 10% fetal bovine serum. Expression vectors were transduced into cells using Lipofectamine 2000 (Invitrogen) or FuGENE 6 (Roche). For drug selection, puromycin (1 μg/μl) was added to the medium 18 h after transfection and the cells cultured for another 48 h in selective medium. The cells were then incubated in puromycin-free medium for 24 h before use. As indicated, cells were incubated with 100 μM cycloheximide (CHX) before collection for Western blot analysis.

### Antibodies

For Chromatin immunoprecipitation (ChIP): LDB1 (Santa Cruz, sc-365074), LMO2 (Santa Cruz, sc-65736), FLAG (Cell Signaling, 14793S), HA (Cell Signaling, 3724S), CTCF (Millipore, 07-729), SSBP3 (Invitrogen, PA5-107280), H3K27ac (RevMAb, 31-1056-00), Histone H2Av (Active Motif, 61751).

For Western Blot (WB): HA (Cell Signaling, 3724S), SSBPs (Abcam, ab177944; claimed to recognize SSBP2, but also recognizes SSBP4 based on our experiments (Figure S3C and D), LDB1 (Cell Signaling, 55476S), LMO2 (Santa Cruz, sc-65736), beta ACTIN (Santa Cruz, sc-47778).

The SSBP3 antibody used for *in vitro* dimerization assays and EMSA has been described ^59^. Immunoglobulin G (IgG, sc-2027) and antibodies to LHX2 (sc-19342X) and LDB1 (sc-11198X) were purchased from Santa Cruz Biotechnology.

### SSBP2-eGFP-mAID-2xFLAG knock-in cell line

A DNA fragment containing LHA (1180 bp left homology arm)-eGFP-mAID-2xFLAG-RHA (1276 bp right homology arm) were cloned into Zero Blunt TOPO vector (Invitrogen, 450245), amplified by Phusion (NEB, M0531S) and purified with the QIAquick PCR Purification Kit (QIAGEN, 21804). sgRNA targeting *Ssbp2* stop codon (GGCAACGGATCACACGCTCA) was cloned into pX330-mCherry vector (Addgene, 98750) using BbsI (NEB, R0539S). Maxi prep was performed with the PureLink HiPure Plasmid Maxiprep Kit (Invitrogen, K210006). 6 ug PCR product and 18 ug pX330-mCherry plasmid were electroporated into G1E-ER4 cells using the Amaxa II electroporator (Lonza, program G-016) with the Amax II Cell Line Nucleofector Kit (R) (Lonza, VCA-1001). mCherry+ cells were sorted by fluorescence activated cell sorting at 48 h post-transfection, followed by single-cell clone screening by PCR and further confirmed by Sanger sequencing.

### 2xFLAG-SSBP4 knock-in cell line

A DNA fragment containing LHA (1002 bp left homology arm)-2xFLAG-RHA (1068 bp right homology arm) was cloned into Zero Blunt TOPO vector (Invitrogen, 450245). sgRNA targeting *Ssbp4* start codon (TGACAGGTGCGGAGCATGTA) was cloned into pX330-mCherry vector (Addgene, 98750) using BbsI (NEB, R0539S).

### SSBP2/4 double knock out cell lines

We transduced G1E-ER4 cells with AsCas12a^67^ lentivirus made from pRG232 (Addgene, 149723). sgRNA synthetic oligonucleotides targeting SSBP2 (TGAGCTCCTACATGGAGCAGATA), SSBP3 (TACGTCTACGAATATTTACTGCA), and SSBP4 (GCCTCACTGGAGTGTTCACAGGC) LUFS domains (Eurofins Genomics) were subcloned into lentiviral vector pRG212 (Addgene, 149722). A second transduction was performed with lentivirus made from pRG212 containing the three SSBPs guides. GFP+ cells were sorted by fluorescence activated cell sorting at 48 h post-transfection, followed by single-cell clone screening by PCR and further confirmed by Sanger sequencing.

### SSBP3-FKBP12^F36V^-2xHA knock-in cell line in *SSBP2/4* double knock out cells

A DNA fragment containing LHA (1310 bp left homology arm)-FKBP12^F36V^-2xHA -RHA (1277 bp right homology arm) was first cloned into Zero Blunt TOPO vector (Invitrogen, 450245). Then it was amplified by Phusion (NEB, M0531S) and purified by QIAquick PCR Purification Kit (QIAGEN, 21804). sgRNA targeting *SSBP3* stop codon (TGTGACCCCCTCCGAGACGC) was cloned into pX330-mCherry vector (Addgene, 98750) using BbsI (NEB, R0539S). mCherry+ cells were sorted by fluorescence activated cell sorting at 48 h post-transfection, followed by single-cell clone screening by PCR and further confirmed by Sanger sequencing.

### Western Blotting

Western blotting was performed following standard protocols. Briefly, cell pellets were boiled at 1x Protein Sample Loading Buffer (LICOR, 928-40004). Equivalent amounts of protein (10–20 µg) were loaded onto 4–20% Mini-PROTEAN TGX Precast Protein Gels (Bio-Rad) and separated in 1x SDS running buffer. Separated proteins were transferred onto PVDF membranes and incubated with primary antibodies overnight. IRDye 800CW Donkey anti-Mouse IgG (LICOR, 926-32212) and IRDye 800CW Donkey anti-Rabbit IgG (LICOR, 926-32213) were used as secondary antibodies.

### RNA isolation and RT-qPCR

Total RNA was extracted with RNeasy Mini Kit (QIAGEN, 74104) following manufacturer’s instructions. Genomic DNA was removed by in-column DNase I digestion (Qiagen, 79256). 1000 ng of RNA was converted to complementary DNA using PrimeScript RT Reagent Kit (TAKARA, RR037A). qRT-PCR was performed using a 10 μl reaction volume containing 10 ng cDNA, 1 μl of 10 μM forward and reverse primers, and 5 μl of Power SYBR Green PCR Master Mix (Thermo Fisher Scientific, 4367660). The reaction volume was adjusted to 10 μl with nuclease-free water. qRT-PCR was conducted on an ABI Vii7 Real-Time PCR System. The ΔΔ^Ct^method (ΔΔCt = ΔCt (treated sample) – ΔCt (control), ΔCt (treated sample or control) = Ct (gene of interest) – Ct (endogenous control)) was used for quantifications. Values of *Gapdh* or β*-Actin* were used as the endogenous control as indicated.

### 4C-Sequencing

To generate 4C templates, 10 million cells were fixed and processed as previously described^80^. DpnII and NlaIII were used as first and second restriction enzyme, respectively. The following 4C primers were used to amplify viewpoints of interest, using 200 ng 4C template input in 4 independent PCR reactions (4 x 50 μl): β*-major* promoter TACACGACGCTCTTCCGATCTGCAGAGCATATAAGGTGAGGTAGGATC and ACTGGAGTTCAGACGTGTGCTCTTCCGATCTCTTTTCTTTGTTTCTCAGTTTGAGTGCATG Pooled reactions were purified by QIAquick PCR Purification Kit (QIAGEN, 21804) eluting in 50 μl milliQ water. 10 μl purified PCR product was used as input material for a second PCR to obtain full Illumina adapter sequences using a universal forward primer (AATGATACGGCGACCACCGAGATCTACACTCTTTCCCTACACGACGCTCTTCCGATCT) and a barcoded reverse primer (CAAGCAGAAGACGGCATACGAGATXXXXXXGTGACTGGAGTTCAGACGTGTGCT) with a 6-nucleotide barcode (XXXXXX). Pooled 4C libraries were purified with Ampure XP beads (Beckman Coulter, A63881) before sequencing on the Illumina NextSeq 2000 platform. 4C sequencing data analysis was performed by pipe4C (https://github.com/deLaatLab/pipe4C).

### ChIP-seq and data analysis

Dual crosslinking was performed at RT for 30 min with 2 mM disuccinimidyl glutarate (DSG) in DPBS followed by addition of formaldehyde to a final concentration of 1% for 15 min. Crosslinking was quenched with 150 mM glycine for 5 min at RT. 10 m dual crosslinked cells were lysed in 1 ml ice-cold cell lysis buffer (10 mM Tris pH 8.0, 10 mM NaCl, 0.2% Igepal, 10 ul/ml PMSF, 2 ul/ml Protease Inhibitors) for 20 min. Nuclei were pelleted and lysed using 1 mL Nuclear Lysis Buffer (50 mM Tris pH 8.0, 10 mM EDTA, 1% SDS, 10 ul/ml PMSF, 2 ul/ml Protease Inhibitors) for 20 min on ice. Samples were sonicated with the Bioruptor Pico (Diagenode) for 15 min: 30sec on, 30sec off, ‘medium’ mode. Sonicated extracts were diluted 10 times with IP Dilution Buffer (20 mM Tris pH 8.0, 2 mM EDTA, 150 mM NaCl, 1% Triton X-100, 0.01% SDS, 10 ul/ml PMSF, 2 ul/ml Protease Inhibitors). For ChIP that needed Drosophila Schneider 2 (S2) spike-in chromatin to normalize genome wide changes, add 2∼5% homemade S2 chromatin or commercial S2 chromatin (Active Motif, 53038) to the diluted chromatin before IP. 100 uL diluted chromatin was taken as input.

For each ChIP, 100 μl Protein G Dynabead (Invitrogen, 10003D) slurry and 10 μg of antibody was used. Chromatin incubation with antibodies was carried out at 4°C with end-to-end rotation for at least 6 h. After IP, beads were washed 4 times with wash buffer (50 mM HEPES KOH pH 7.5, 500 mM LiCl, 1 mM EDTA, 1% NP-40, 0.7% Na-deoxycholate, 10 ul/ml PMSF, 2 ul/ml Protease Inhibitors) and followed by one more wash in TE buffer. Chromatin was then eluted in 200 ul elution buffer (50 mM Tris HCl pH 8.0, 10 mM EDTA, 1% SDS) at 65°C with shaking. 8 ul RNase A (10mg/ml) was added to input and eluted chromatin and incubated at 37°C for 30 min. 10 ul 20mg/ml proteinase K was then added, and subsequent reverse crosslinking was performed at 65°C overnight. DNA was purified using QiAquick PCR purification kit (QIAGEN, 28104). ChIP-qPCR was performed using a Power SYBR Green kit (Invitrogen, 4368577) with signals detected by a ViiA7 System (Life Technologies). ChIP-seq libraries were prepared using NEBNext Ultra II DNA Library Prep Kit (NEB, E7645S) with NEBNext unique dual index primer pairs (NEB, E6440S). Library purification was performed by using AMPure XP Bead (Beckman Coulter, A63881) on a magnetic strip. Libraries QC and sequencing (2x50bp) were performed on an Illumina NextSeq 2000 platform.

ChIP-seq QC and mapping were performed by the nf-core/chipseq pipleline (version 2.0.0)^81^ with the default setting using mm9 genome build. MACS2 (2.2.7.1)^82^ using q ≤ 0.01 to call narrow peaks for LDB1, LMO2, SSBP2/3/4, CTCF. MACS2 (2.2.7.1)^82^ using q ≤ 0.01 to call broad peaks for H3K27ac. For samples not involved in differential analysis, the corresponding bigwig files were simply normalized by RPKM.

DESeq2 (1.44.0)^83^ was used for differential binding analysis between undifferentiated and differentiated conditions; wide type cells and SSBP2/4 double knockout cells. For samples normalized by DESeq2, all the corresponding bigwig files were generated by deeptools (3.5.5)^84^ bamCoverage using the size factors generated by DESeq2 (1.44.0)^83^.

For ChIP-seq supplemented with *Drosophila* Schneider 2 (S2) spike-in chromatin, the reads were mapped separately using mm9 and bdgp6. Normalization factor was computed by using 1M divided by reads mapped to *Drosophila* according to published method^85^. For samples with S2 spike-in normalized, all the corresponding bigwig files were generated by deeptools (3.5.5)^84^ bamCoverage using the normalization factors based on S2 spike-in.

### RNA-seq data analysis

RNA seq library preparation and sequencing were performed as described^25^. RNA-seq analysis were performed by the nf-core/rnaseq pipleline (version 3.14.0)^81^

### PRO-seq

PRO-seq^77^ was performed based on a fast PRO-seq version^86^. For each library, 10 million cells were used together with 0.3 million Drosophila Schneider 2 (S2) cells added as spike-in to control for potential bias associated with library scaling. After 1h DMSO and dTAG^V^-1 treatment, cells were collected simultaneously and processed in pairs. Run-on experiments were performed by adding 2x nuclear run-on (NRO) buffer as described in the original protocol. Two biotin-NTPs (biotin-11-CTP, biotin-11-UTP) were supplied at equal ratio to non-labeled ATP and GTP. To facilitate the removal of PCR duplicates, we added random hexamers to the 5′ end of the REV3 adaptor and 3’ end of REV5 adaptor as unique molecular identifiers (UMI). Reads with the same UMI were collapsed into one. Size-selected libraries were pooled and sequenced on an Illumina Novaseq X plus platform at Novogene or on an Illumina NextSeq 2000 platform.

Fastq sequencing files were processed by fastp^87^ to remove adaptors and low-quality reads and bases; UMI-tools^88^ was used to extract UMI from reads; BWA-MEM^89^ was used for reads mapping using mm9 genome build; SAMtools^90^ was used to convert SAM files to BAM files and BAM file indexing; UMI-tools^88^ was used again for deduplication based on the extracted UMI; RPKM was used for normalization to generate bigwig from deeptools bamCoverage^84^. Quantification of PRO-seq signals for each gene from transcription start site (TSS) to transcription end site (TES) and differential gene expression analysis was achieved by HOMER (getDiffExpression.pl) (http://homer.ucsd.edu/homer/ngs/peaksReplicates.html).

### Micro-C

Micro-C was performed as previously described^70^ with minor adjustments. 5 million cells were crosslinked with 1% formaldehyde for 10 min and quenched by 125 mM glycine for 5 min, followed by an additional fixation with 3mM DSG (ProteoChem, c1104-1gm) for 40 min and 5 min 0.4M glycine quenching. Fixed cells were permeabilized with Micro-C Buffer 1 at a concentration of 1 million cells/100uL (50 mM NaCl, 50 mM Tric-HCl pH 7.5, 5 mM MgCl_2_, 1 mM CaCl_2_, 0.2% NP-40, 1x Protease Inhibitor Cocktail tablet (Millipore Sigma, 11836170001)) for 20 min on ice. 1,000 gel units Micrococcal nuclease (NEB, M0247S) was added to the permeabilized nuclei for 20 min at 37°C with rotation (850 rpm) to digest chromatin to 90% monomer/10% dimer ratio. The exact units of micrococcal nuclease were tested each time before proceeding to digest actual samples. EGTA was added at a final concentration of 4 mM to stop reaction for 10 min at 65°C. Nuclei were washed twice with Micro-C Buffer 2 at a concentration of 1 million cells/100uL (50 mM NaCl, 50 mM Tric-HCl pH 7.5, 10 mM MgCl_2_, 100ug/ml BSA, 1x Protease Inhibitor Cocktail tablet (Millipore Sigma, 11836170001)) and then resuspended in 50 uL 1X NEBuffer 2.1.

Digested fragments were de-phosphorylated with 5 U r-SAP (NEB, M0371S) for 45 min at 37°C in de-phosphorylation buffer (50mM NaCl, 10mM Tris-HCl, 10mM MgCl_2_, 100 ug/mL BSA). De-phosphorylated fragments were subjected to end-chewing using 40 U large Klenow Fragment (NEB, M0210S) for 15 min at 37°C in the following buffer: 50mM NaCl, 10 mM Tris-HCl, 10 mM MgCl_2_, 100 ug/mL BSA, 2 mM ATP, and 3 mM DTT. 20 U T4 PNK (NEB, M0201S) was added at the end-chewing step. End labeling was achieved by adding biotin-dATP (Jena Bioscience, NU-835-BIO14-S), biotin-dCTP (Jena Bioscience, NU-809-BIOX-S), dTTP, and dGTP and incubating at 25°C for 45 min. Finally, fragmented and labeled DNA ends were ligated using 1,200 U of T4 DNA ligase (NEB, M0202S) and incubating at room temperature overnight. Unligated ends were removed by exonuclease III for 10 min at 37°C. After reverse-crosslinking, di-nucleosomal fragments were collected by AMPure XP Bead (Beckman Coulter, A63881). The recovered di-nucleosomal fragments were immobilized on MyONE Strptavidin C1 Dynabeads (Thermo Fisher, 65001). Micro-C libraries were prepared using NEBNext Ultra II DNA Library Prep Kit (NEB, E7645S) with NEBNext unique dual index primer pairs (NEB, E6440S). Shallow sequencing was done by 2x50bp on an Illumina Nextseq 2000. Deep sequencing was done by 2x150bp on the Illumina Novaseq X plus at Novogene.

### Micro-C data processing and visualization

Distiller pipeline (v3.3) (https://github.com/mirnylab/distiller-nf) was used to generate contact maps from fastq files. PCR duplicates were removed from each replicate and balanced contact maps were generated for each treatment condition from merged biological replicates. Iterative correction and eigenvector decomposition (ICE) balancing^91^ was used to normalize contact maps using default settings. Log-binned contact decay curves (a.k.a. P(s) curves) were generated for contact maps from each biological replicate using cooltools (v0.5.1)^92^. We used coolbox (v0.3.8)^93^ to visualize contact maps and aligned ChIP-seq, PRO-seq tracks.

### Micro-C compartment analysis

We used cooltools (v0.5.3)^92^ to compute cis eigenvector 1 values based on 100kbp resolution merged contact maps as well as maps from each biological replicate from DMSO and dTAG^V^-1 treated conditions. Saddle plots showed consistent A-to-B or B-to-A switches in all biological replicates.

### Micro-C domain analysis

We identified domains using the rGMAP (v1.4)^94^ software using 10kb-binned contact matrices. We called domains for each 10kbp contact map with two iterations. For the first domain sweep, we used default settings. For the second domain sweep, we used parameters dom_order=3, maxDistInBin=50 to capture smaller domains that might be missed during the first iteration.

Based on these preliminary domain calls, we calculated insulation scores at associated preliminary domain boundaries using a 120kbp sliding diamond with cooltools. We fine-tuned boundary positions based on minimum insulation score within a +/-60kb window of the preliminary boundaries. Domains overlapping sliding windows with sum counts ≤12 raw counts were removed due to low coverage.

We next merged domain called from DMSO and dTAG^V^-1 treated samples if they were within −/+80kbp of each other. Low-mappable, unstructured areas, domains that were smaller than 100kb and larger than 2mb were removed.

### Loop calling and quantification

Mustache^95^ was used to identify loops using merged contact maps for each treatment condition. We called loops on 1kb, 2kb, 5kb, and 10kb resolution contact maps separately for each treatment condition using the default settings. We created a master loop list by merging DMSO and dTAG^V^-1 treated loops from each resolution. Redundant loops were merged (redundancy defined as being the same pixel or adjacent pixels). Then, we merged consensus lists from each resolution (1kb, 2kb, 5kb, and 10kb), retaining the smallest resolution coordinates in instances where a loop was called at multiple resolutions. To quantify loops, we used the approach outlined in Lam et al.^96^. Briefly, we quantified loop strength as an observed over locally-adjusted expected value. This approach considers enrichment of a loop relative to its local ‘donut’ shaped surroundings. For a given loop’s peak pixel, the observed value was first calculated at the resolution at which the loop was called. A locally-adjusted expected value was calculated by multiplying the global expected value at the loop’s peak pixel by a local-adjustment value (i.e. the sum of the observed contacts within the rounded donut region divided by the sum of the expected contacts within the rounded donut). For each loop, loop strength was calculated in both DMSO and dTAG^V^-1 conditions. To quantify loop changes after auxin treatment, we calculated log2(dTAG^V^-1 loop strength / DMSO loop strength). We annotated looping change based on a 1.5-fold change threshold. Loops with change ≥log2(1.5) as “strengthened”, change ≤ −log2(1.5) as “weakened”, and all other loops as “unchanged.

### Plasmid constructions

Expression plasmids pTracer-EF/V5-HisLDB1 and pcDNA3.1(+)-LDB1 have been described^55^. For *in vitro* transcription-translation reactions, PCR products were subcloned into pcDNA3.1(+) to generate, respectively, pcDNA3.1(+)-SSBP3, -LDB1(50-375) and -LDB1(200-375). To construct pcDNA3.1(+)-TD-LDB1, pcDNA3.1(+)-LDB1 was used as a template, the LDB1 open reading frame was amplified, fused at its 3’ end with a linker encoding 22 glycines (forward oligonucleotide AAGAATTCATGCTGGATCGGGATG and reverse oligonucleotide TGTACCTCCGCCGGACCCTCCGCCGCTCCCACCTCCAGAACCCCCGCCGGATCCACCCCCGGTTCCTCCCTGGGAAGCCTGTGACGTGGG), and then further amplified to generate *EcoRI* and *EcoRV* restriction sites at its ends (forward primer AAGAATTCATGCTGGATCGGGATG and reverse primer AAGATATCTGTACCTCCGCCGGACCC). *EcoRI*- and *EcoRV*-digested PCR product was subcloned into pTracer-EF/V5-HisLDB1 to construct pTracer-EF/V5-HisTD-LDB1. Finally, TD-LDB1 was transferred into pcDNA3.1(+) to create pcDNA3.1(+)-TD-LDB1. The correct sequences of their inserts and the sizes of the proteins expressed from these plasmids were verified.

Site-Directed Mutagenesis: The third nucleotide upstream of the start codon in vector pcDNA3.1(+)-TD-LDB1 was changed from T to G using site-specific mutagenesis, CCAGTGTGGTGGAA**G**TC<u>ATG</u>CTGGATCGGGCTC to enhance translation. The sequence of the mutated DNA construct was confirmed by nucleotide sequencing and its ability to direct expression of a protein of the predicted size was verified by Western blot analysis.

### *In vitro* transcription-translation and chemical crosslinking

^35^S-labeled proteins were synthesized using the TNT T7 Coupled Reticulocyte Lysate System (Promega) and translation-grade ^35^S-Met (GE Healthcare). 8-12 µl of the translation reaction mixture containing ^35^S-methionine were added to 100 µl of 50 mM HEPES (pH 7.4), 150 mM NaCl, 1% Triton X-100, 10% glycerol and protease inhibitors. Following centrifugation at 14,000 *g* for 30 min at 4°C, freshly prepared bis (succinimidyl) suberate (BS3, Thermo Scientific) was added at a final concentration of 3 mM, the mixture incubated for 30 min at room temperature, and the reaction terminated after an additional 15 min with addition of 250 mM glycine prior to SDS-PAGE and fluorography.

### Electrophoretic mobility shift analysis

Nuclear extracts from αT3-1 cells obtained from Dr. Pamela Mellon (University of California, San Diego) were prepared and electrophoretic mobility shift analysis (EMSA) carried out as detailed^59^. SSBP3 antibody has been described previously^59^. Immunoglobulin G (IgG, sc-2027) and antibodies to LHX2 (sc-19342X) and LDB1 (sc-11198X) were purchased from Santa Cruz Biotechnology. Protein-DNA complexes were fractionated by electrophoresis in 4% polyacrylamide gels in Tris-glycine buffer for 16 h at 4°C and the dried gels subjected to autoradiography. A probe containing a minimal LHX2 binding element with the sequence ATATCAGGTACTTAGCTAATTAAATGT was used.

### DNA-linking assay

This assay was described previously in detail^66^. Briefly, 5 µg of nuclear protein extract were incubated with 2 nM ^32^P-labeled double-stranded probe at room temperature for 5 min, 5’-biotinylated double-stranded probe added at a concentration of 4 nM, and the reaction allowed to continue for an additional 15 min. As noted, 2 µg of anti-biotin antibody, protein A (Sigma) and/or 2.5 U of streptavidin-conjugated β-galactosidase (Roche) were added and the mixture incubated for 10 min. Protein-DNA complexes were resolved and visualized as described above for standard EMSA.

### Autoradiographic analysis

Band intensities on autoradiographs were quantified with the ImageJ software program (National Institutes of Health; http://rsb.info.nih.gov).

## ACKNOWLEDGMENTS

The authors thank members of the Blobel lab for helpful discussions and reading of the manuscript. Flowcytometry was performed at the flow core of the Children’s Hospital of Philadelphia. The sequencing was performed in the Huck Genomics Research Incubator RRID:SCR_024530 at Penn State University. This work was supported by grant R01DK05937 to G.A.B.; T32GM008216, 1F31DK136200-01A1, and the Blavatnik Family Fellowship Award to N.G.A.; T32HG000046 and F30DK132824 to J.C.L.; R24DK106766 to R.C.H. and G.A.B.

## AUTHOR CONTRIBUTIONS

X.W., Y.C., S.J.B. and G.A.B. conceived the study. X.W., Y.C., K.A.H., J.X. performed the experiments. X.W. analyzed the next generation sequencing data. N.G.A. provided LDB1-AID cell line and shared LDB1 Micro-C, TT-seq data. J.C.L and N.G.A shared their protocols to analyze Micro-C data. L.N. provided SSBP3 antibody for protein crosslinking and EMSA assays. Next generation sequencing was performed by C.A.K. and B.M.G. under supervision of R.C.H. The manuscript was written by X.W., Y.C., S.J.B. and G.A.B. with input from all authors.

## Figure Legends

**Figure S1.**
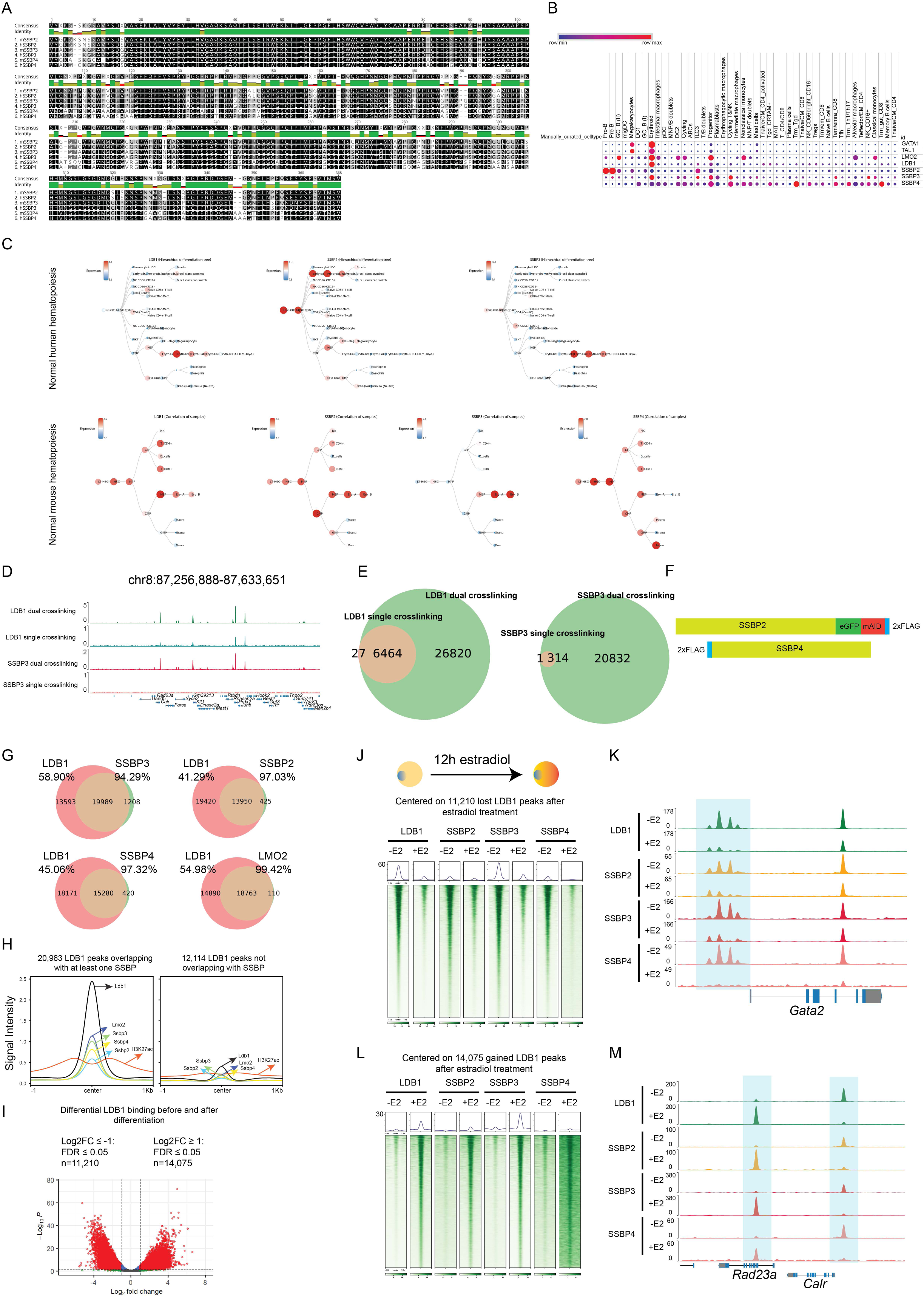
SSBP2/3/4 and LDB1 colocalized in both undifferentiated and differentiated conditions, related to Figure 1. (A) Protein sequence alignments of human and mouse SSBP2/3/4, generated by Geneious software. (B) SSBP2/3/4 expression across diverse human blood cell types. (C) SSBP2/3/4 expression in normal human and mouse hematopoiesis. (D) ChIP-seq tracks showing dual crosslinked LDB1 (n=5), SSBP3 (n=2) and single crosslinked LDB1 (n=2), SSBP3 (n=2). (E) Left: venn diagram showing the overlapping and unique peaks between dual crosslinked LDB1 (n=5) and single crosslinked LDB1 (n=2); right: venn diagram showing the overlapping and unique peaks between dual crosslinked SSBP3 (n=2) and single crosslinked SSBP3 (n=2). (F) Upper: Diagram showing eGFP-mAID-2xFLAG was knocked in at the C terminus of SSBP2; Lower: Diagram showing 2xFLAG was knocked in at the N terminus of SSBP4. (G) Venn diagram showing the overlapping and unique peaks between LDB1 (n=5) and SSBP3 (n=2), SSBP2 (n=5), SSBP4 (n=4), LMO2 (n=3). (H) Metaplots of LDB1 (n=5), LMO2 (n=3), SSBP2 (n=5), SSBP3 (n=2), SSBP4 (n=4), and H3K27ac (n=2) on LDB1 peaks overlapping with at least one SSBP (left) or LDB1 peaks not overlapping with any SSBP (right). (I) Volcano plot showing differential LDB1 binding before (n=5) and after differentiation (n=3). (J) Upper: Diagram showing the differentiation process. Lower: heatmaps centered on lost LDB1 peaks showing LDB1(-E2: n=5; +E2: n=3), SSBP2 (-E2: n=5; +E2: n=2), SSBP3 (-E2: n=2; +E2: n=2), and SSBP4 (-E2: n=4; +E2: n=2) binding before and after differentiation. E2 represents estradiol. (K) ChIP-seq tracks showing LDB1(-E2: n=5; +E2: n=3), SSBP2 (-E2: n=5; +E2: n=2), SSBP3 (-E2: n=2; +E2: n=2), and SSBP4 (-E2: n=4; +E2: n=2) binding before and after differentiation at the *Gata2* locus. E2 represents estradiol. (L) Same as (J) but for gained LDB1 peaks. (M) Same as (K) but for *Rad23a* locus.

**Figure S2.**
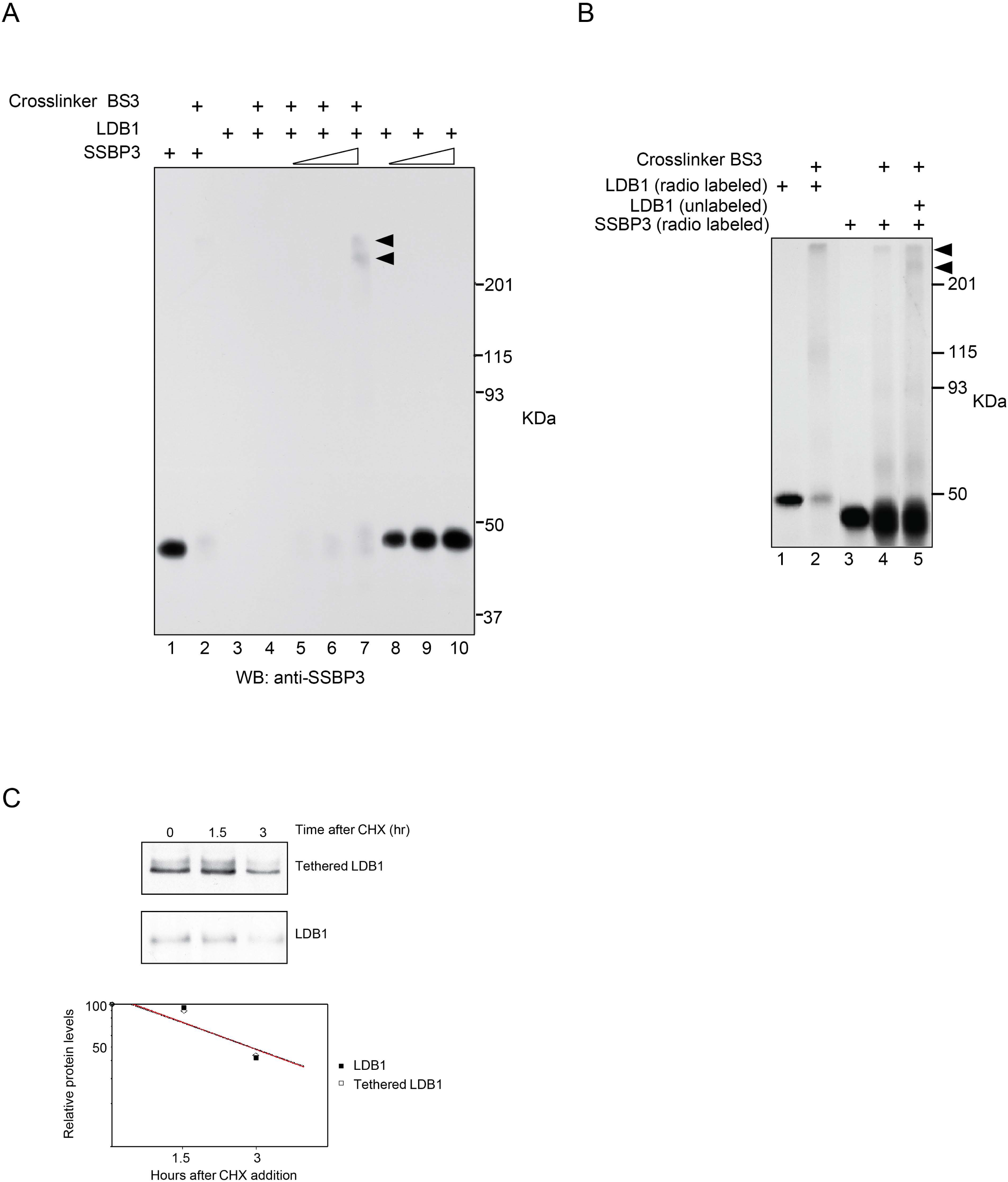
SSBP3 forms complex with LDB1 dimer, related to Figure 2. (A) Western blot using anti-SSBP3 antibody analyzes products from crosslinking of SSBP3 and LDB1 *in vitro*. (B) Autoradiograph of proteins separated by SDS-PAGE after crosslinking using radio labeled SSBP3 and unlabeled “cold” LDB1. High molecular weight complexes containing LDB1 and SSBP3 are shown by solid arrowheads. (C) Western blot quantification of protein turnover of endogenous LDB1 and transfected tethered LDB1 in 293T cells. Cells were transfected with tethered LDB1 and treated with 100 µM cycloheximide (CHX) for the times indicated (Top two panels). Comparison of densitometry readings of X-ray films (Bottom panel).

**Figure S3.**
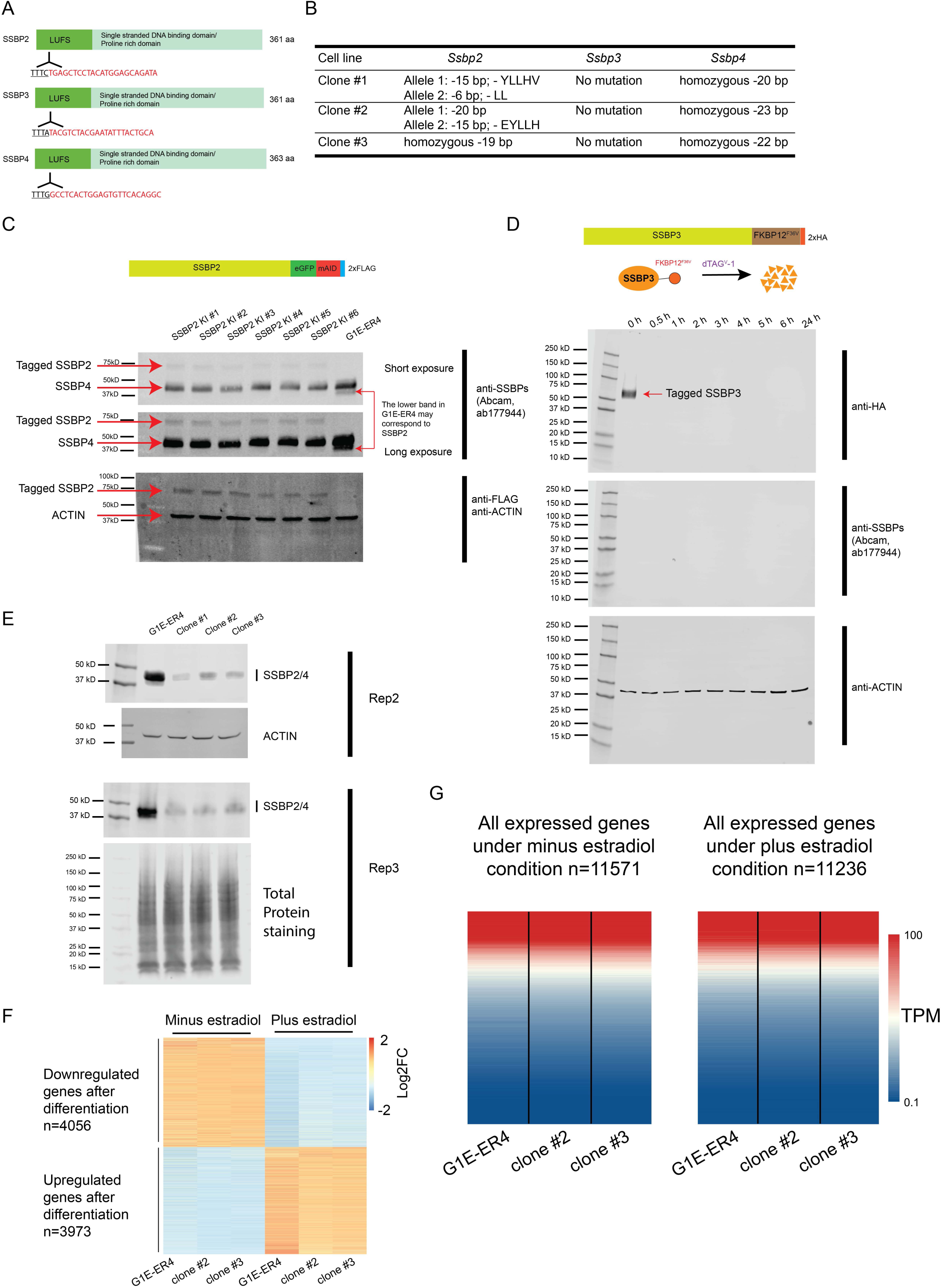
Dual loss of SSBP2 and SSBP4 minimally affected gene expression, related to Figure 3. (A) Diagram showing three guides targeting each of the N terminal LUFS domain of SSBP2/3/4 for CRISPR/Cas12a-mediated gene editing. (B) Mutation details in *Ssbp2/4* hypomorphic (clone #1 and #2) and knockout (clone #3) cells. (C) Western blot analysis of total proteins from SSBP2-eGFP-mAID-2xFLAG and G1E-ER4 cells. FLAG antibody was used to confirm the presence of SSBP2-eGFP-mAID-2xFLAG protein. SSBPs (Abcam, ab177944) was used to show this antibody recognize not only SSBP2 but also SSBP4. ACTIN was used as a loading control for the inputs. WB was performed only once. (D) Western blot analysis of total proteins from SSBP3-FKBP12^F36V^-2xHA cells treated with 500 nM dTAG^V^-1 over a time course. HA antibody was used to confirm the presence and degradation of SSBP3-FKBP12^F36V^-2xHA protein. SSBPs (Abcam, ab177944) was used to show this antibody cannot react with SSBP3. ACTIN was used as a loading control for the inputs. WB was performed twice, and representative result is shown. (E) Western blot with SSBPs antibody (Abcam, ab177944) in G1E-ER4 and *Ssbp2/4* hypomorphic (clone #1 and #2) and knockout (clone #3) cells. ACTIN or total protein staining was used as a loading control. (F) Heatmap showing gene expression levels between G1E-ER4 (-E2: n=1; +E2: n=1) and clone #2 (-E2: n=1; +E2: n=1), clone #3 (-E2: n=1; +E2: n=1). The heatmap was generated based on the differentially expressed genes after 24h estradiol treatment. E2 represents estradiol. (G) Heatmap showing gene expression levels between G1E-ER4 (-E2: n=1; +E2: n=1) and clone #2 (-E2: n=1; +E2: n=1), clone #3 (-E2: n=1; +E2: n=1). The heatmap was generated based on the TPM (transcripts per million) values of all expressed genes under either minus estradiol condition or plus estradiol condition. E2 represents estradiol.

**Figure S4.**
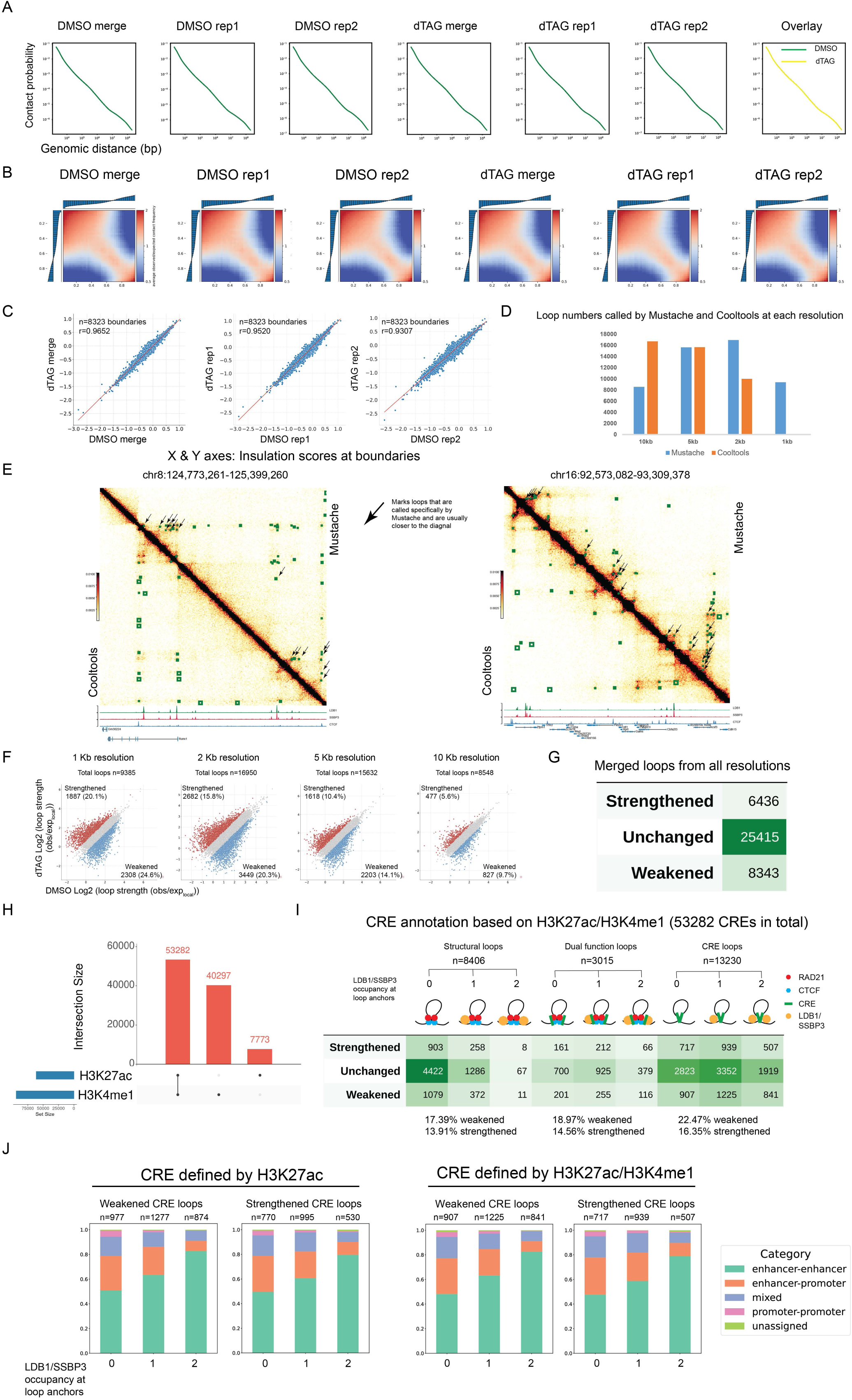
Acute depletion of SSBP3 did not affect compartments and TADs, but affected chromatin loops, related to Figure 3. (A) Micro-C contact-versus-distance curves after 1h DMSO treatment (n=2) or 1h dTAG^V^-1 treatment (n=2). The last lane presents an overlay of contact-versus-distance curves, derived from the replicate-merged DMSO or dTAG Micro-C data. (B) Saddle plots showing whole genome compartment strength after 1h DMSO treatment (n=2) or 1h dTAG^V^-1 treatment (n=2). (C) Scatter plot of Log2 insulation score values from 1h DMSO treatment (n=2) or 1h dTAG^V^-1 treatment (n=2) contact maps for all boundaries called across both conditions. TADs were identified using rGMAP on 10k resolution Micro-C matrices. Insulation scores were calculated using a 120kb sliding window. (D) Comparison of loop numbers called by Mustache and Cooltools at 1kb, 2kb, 5kb, and 10kb resolutions. Cooltools failed to call loops at 1kb resolution. (E) Micro-C contact heatmaps showing pairwise comparison between loops called by Mustache (upper right) and Cooltools (lower left) at two representative loci. LDB1 (n=5), SSBP3 (n=2), and CTCF (n=2) ChIP-seq tracks in G1E-ER4 were added below the heatmaps. Arrows marked selected loops that were called specifically by Mustache, which were usually closer to the diagonal. (F) Scatter plot showing differential looping for loops called at 1kb, 2kb, 5kb, and 10kb resolutions from DMSO treatment or dTAG^V^-1 treatment. (G) Summary of all strengthened, unchanged, and weakened loops merged from all resolutions. (H) Upset plot illustrating the overlapping and unique peaks between H3K27ac and H3K4me1 ChIP-seq data. (I) Upper: Diagram showing three categories of loops: structural loops, dual function loops, and CRE loops, which were further stratified by LDB1/SSBP3 occupancy at loop anchors. Lower: Differential analysis for the indicated loop types after SSBP3 depletion. CRE is annotated based on H3K27ac/H3K4me1. (J) Stacked bar plots illustrate the composition of weakened or strengthened CRE loops (E-E, E-P, P-P) based on LDB1/SSBP3 occupancy at 0, 1, or 2 anchors. CRE is annotated by H3K27ac (left) or H3K27ac/H3K4me1 (right).

**Figure S5.**
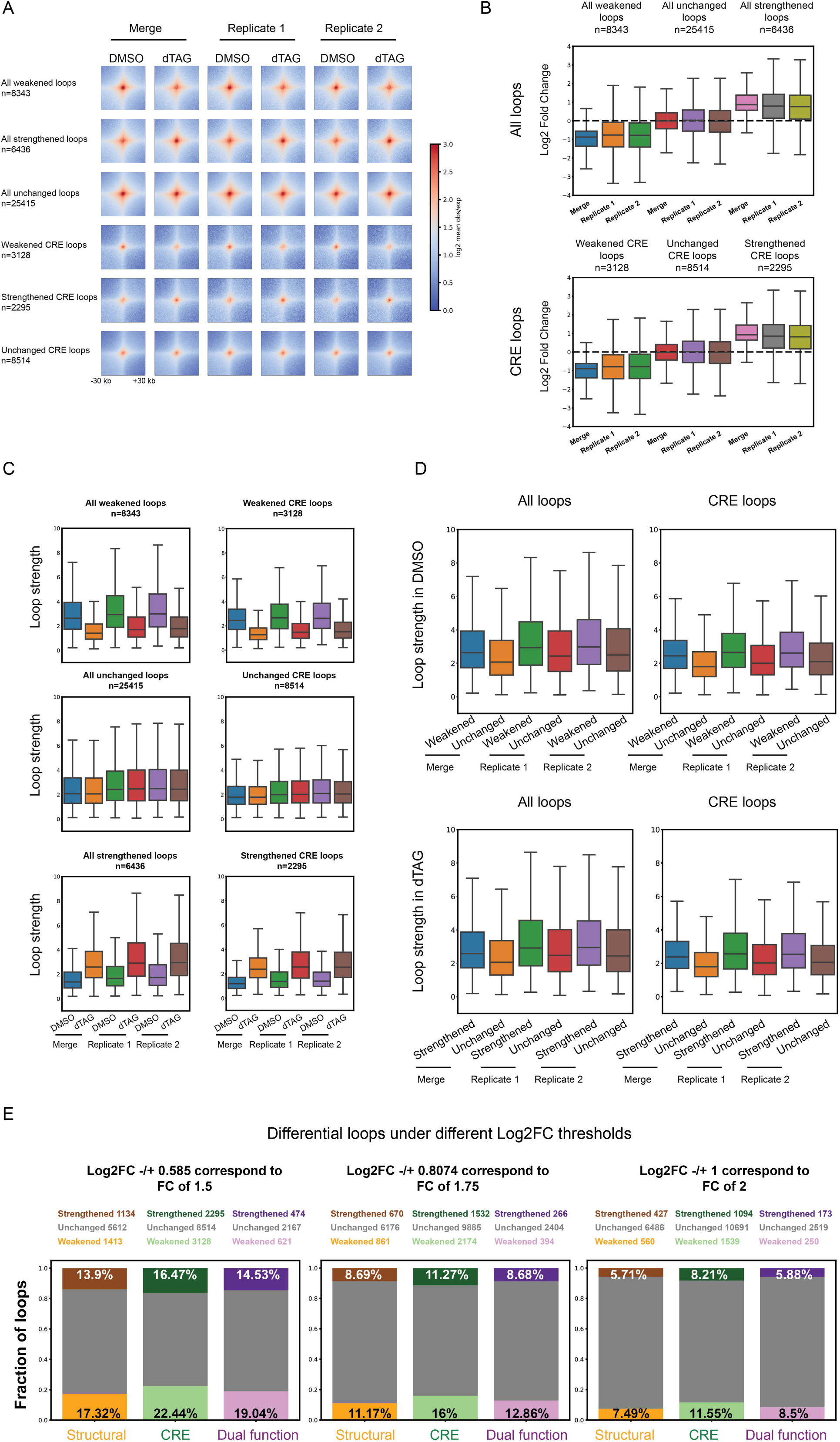
Concordance among replicates for average trends among loop changes, related to Figure 3. Figure 3 (A) Pileup plots for the indicated differential loops in both merged data and individual biological replicates under DMSO and dTAG treatment conditions. (B) Box plots display the Log2 Fold Change for the indicated differential loops following SSBP3 depletion in both merged data and individual biological replicates. (C) Box plots display the loop strength of the indicated differential loops in both merged data and individual biological replicates under DMSO and dTAG treatment conditions. (D) Upper: Box plots compare the loop strength of weakened and unchanged loops in the DMSO treatment condition in both merged data and individual biological replicates. Lower: Box plots compare the loop strength of strengthened and unchanged loops in the dTAG treatment condition in both merged data and individual biological replicates. (E) Stacked bar plots showing the fraction of differential structural, CRE, and dual function loops under different Log2FC thresholds.

**Figure S6.**
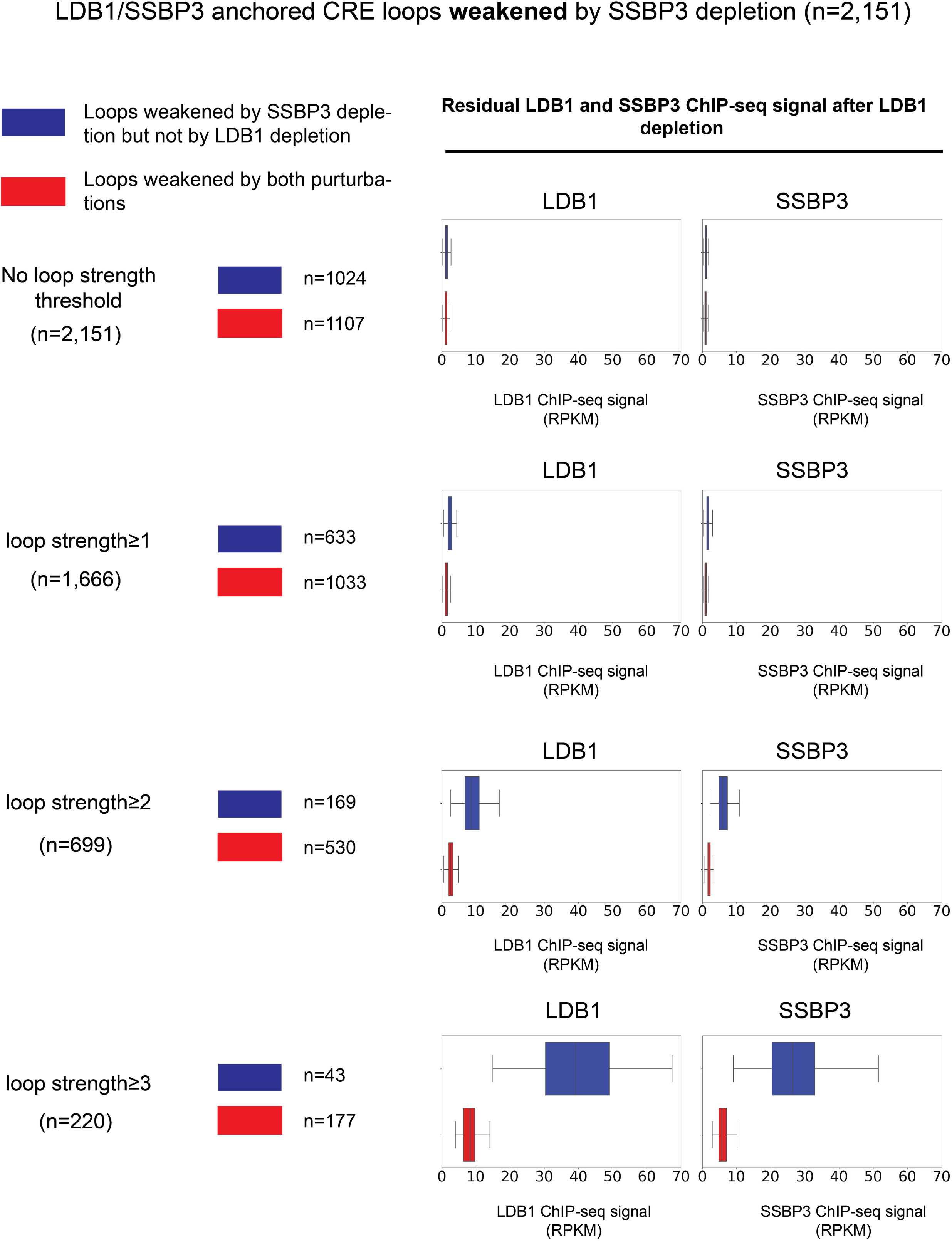
LDB1/SSBP3 anchored CRE loops weakened by SSBP3 depletion were also weakened by LDB1 depletion, related to Figure 3. Boxplots showing residual LDB1 and SSBP3 ChIP-seq signals after LDB1 depletion. The loops that were weakened by SSBP3 depletion but not by LDB1 depletion (blue boxes) had higher residual LDB1 and SSBP3 binding at their loop anchors than loops that were weakened by both SSBP3 depletion and LDB1 depletion (red boxes).

**Figure S7.**
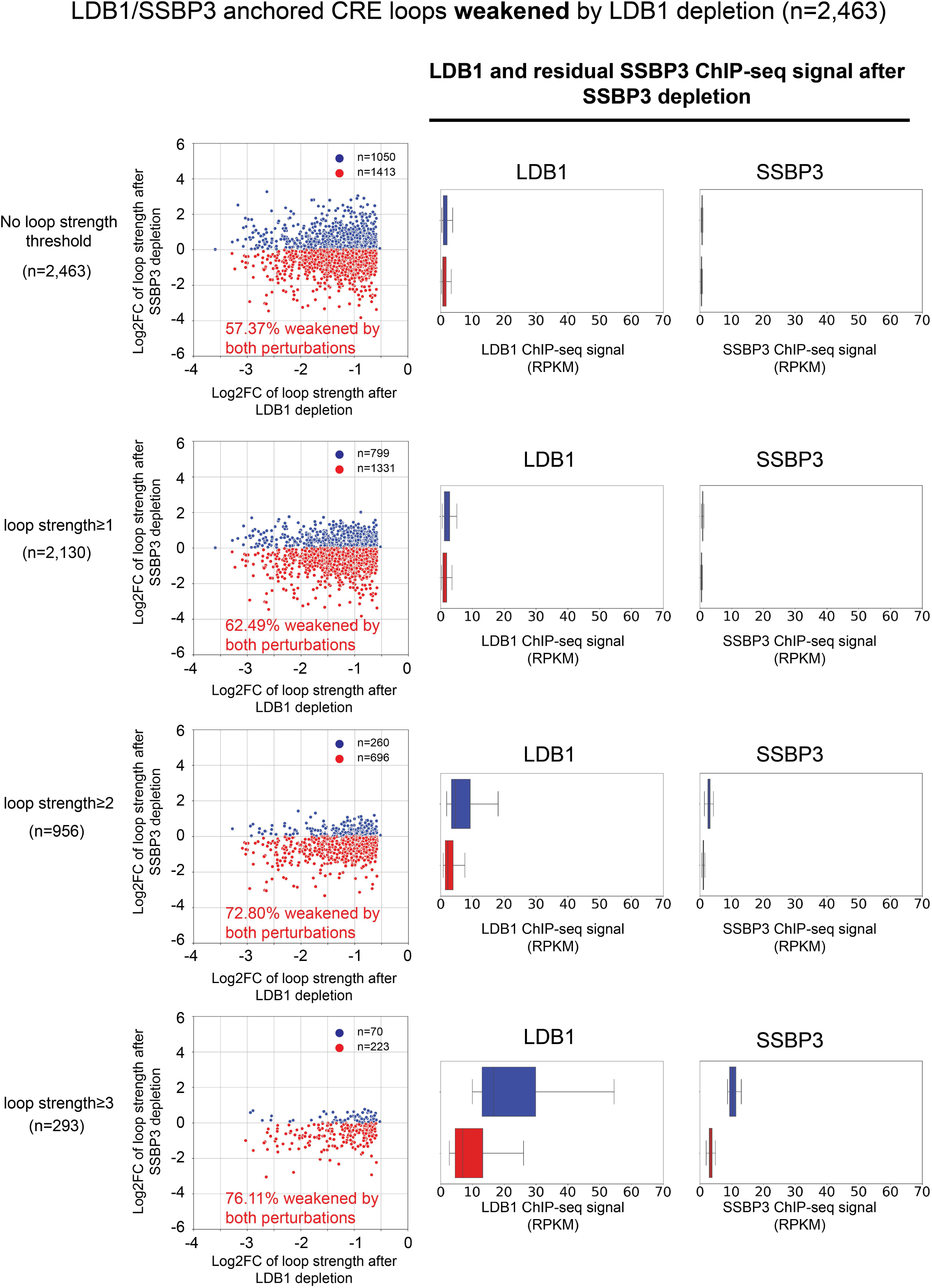
LDB1/SSBP3 anchored CRE loops weakened by LDB1 depletion were also weakened by SSBP3 depletion, related to Figure 3. Left: Scatterplots showing loops that were weakened by both LDB1 depletion and SSBP3 depletion (red dots) and loops that were weakened by LDB1 depletion but not by SSBP3 depletion (blue dots). The more stringent threshold of loop strength (observed over locally adjusted expected value) is used, the higher overlapping rate is observed. Right: Boxplots showing LDB1 and residual SSBP3 ChIP-seq signals after SSBP3 depletion. The loops that were weakened by LDB1 depletion but not by SSBP3 depletion (blue boxes) had higher LDB1 and residual SSBP3 binding at their loop anchors than loops that were weakened by both LDB1 depletion and SSBP3 depletion (red boxes).

**Figure S8.**
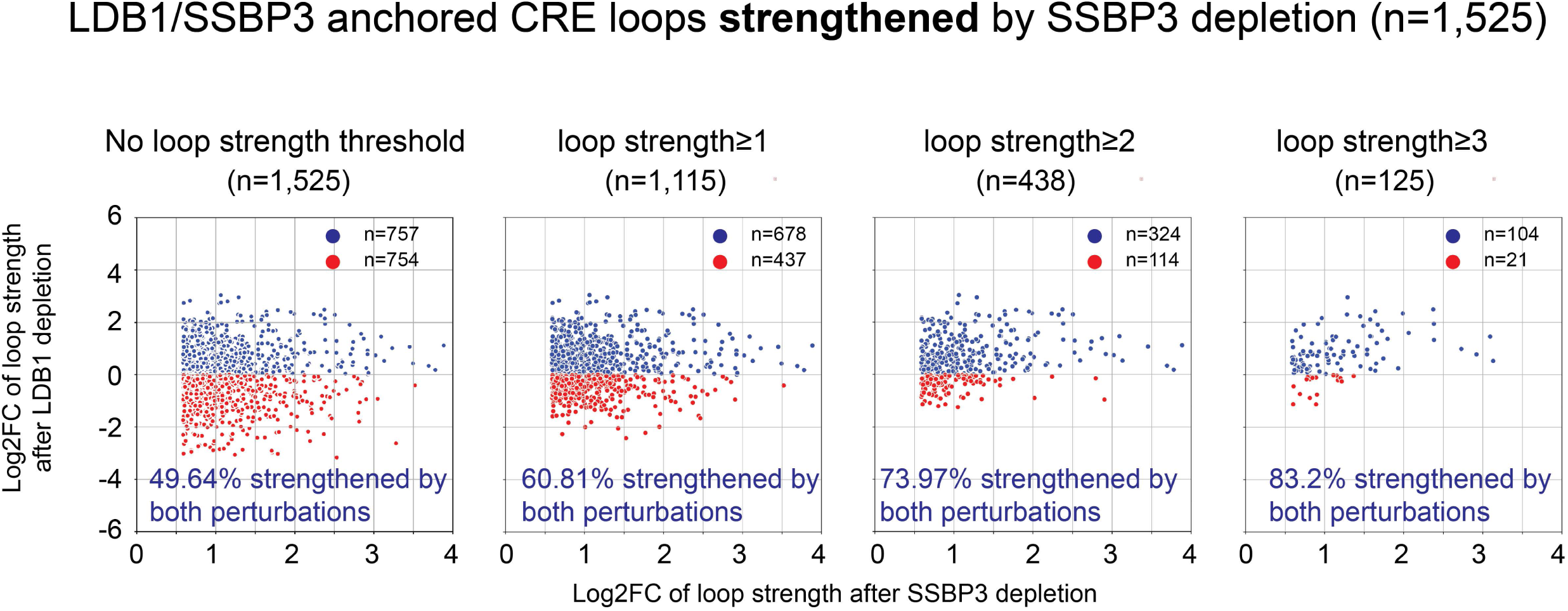
LDB1/SSBP3 anchored CRE loops strengthened by SSBP3 depletion were also strengthened by LDB1 depletion, related to Figure 3. Scatterplots showing loops that were strengthened by both SSBP3 depletion and LDB1 depletion (blue dots) and loops that were strengthened by SSBP3 depletion but not by LDB1 depletion (red dots). The more stringent threshold of loop strength (observed over locally adjusted expected value) is used, the higher overlapping is observed.

**Figure S9.**
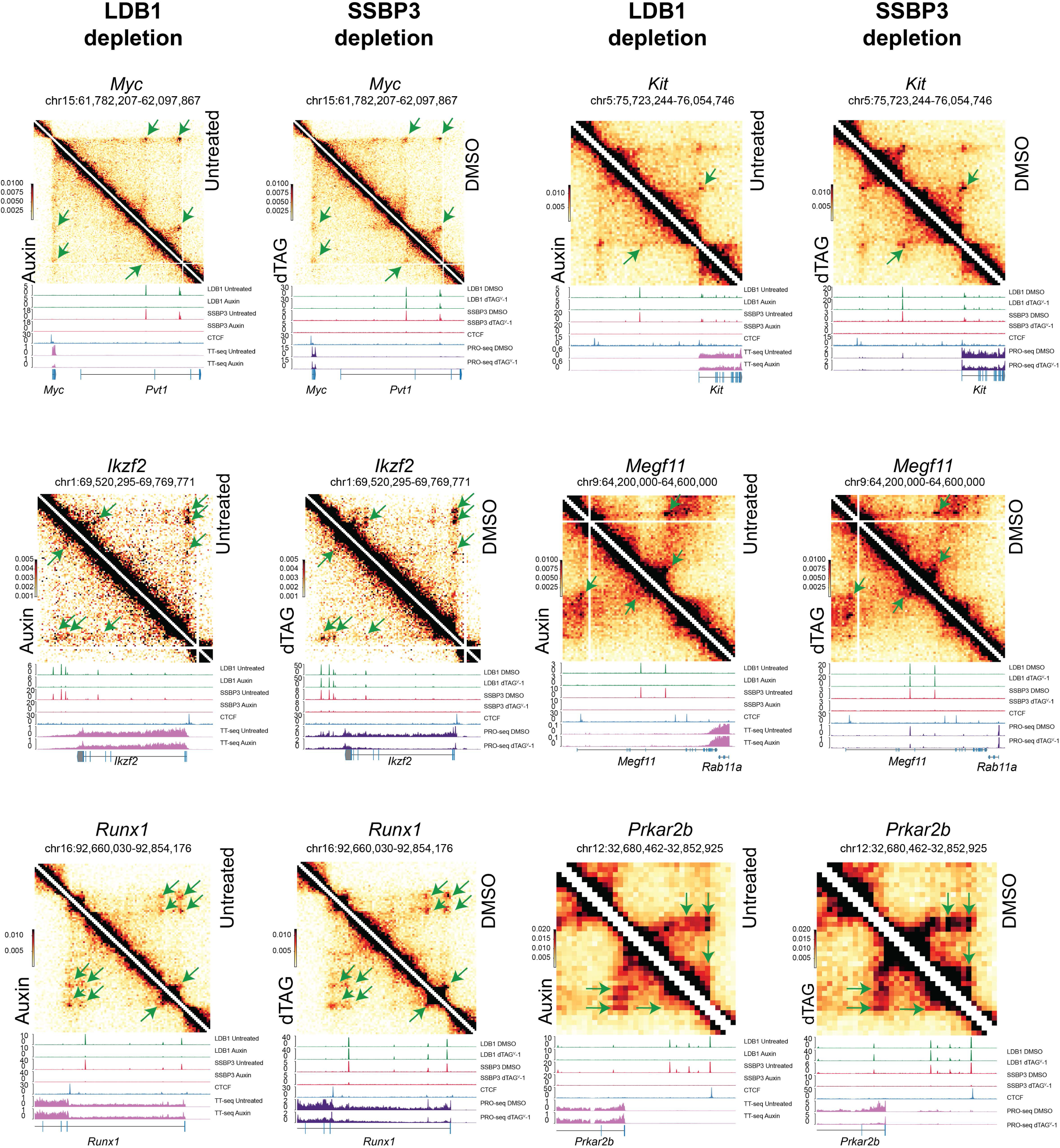
SSBP3 and LDB1 regulate chromatin looping at a same group of loci, related to Figure 3. Pairwise comparisons between Micro-C contact heatmaps after LDB1 depletion and SSBP3 depletion (n=2) at *Myc, Kit, Ikzf2, Megf11, Runx1, and Prkar2b* loci, showing LDB1/SSBP3 anchored loops (marked by green arrows) were weakened by both LDB1 depletion and SSBP3 depletion. Untreated and 4h auxin treated LDB1 (n=2) and SSBP3 (n=2) ChIP-seq tracks were added below the LDB1 Micro-C heatmaps. 1h DMSO treated and 1h dTAG^V-1^ treated LDB1 (n=3) and SSBP3 (n=3) ChIP-seq tracks were added below the SSBP3 Micro-C heatmaps. CTCF ChIP-seq track in G1E-ER4 was added to mark nearby structural loops. TT-seq and PRO-seq data (n=3) were separately incorporated into the Micro-C heatmaps for LDB1 depletion and SSBP3 depletion conditions.

**Figure S10.**
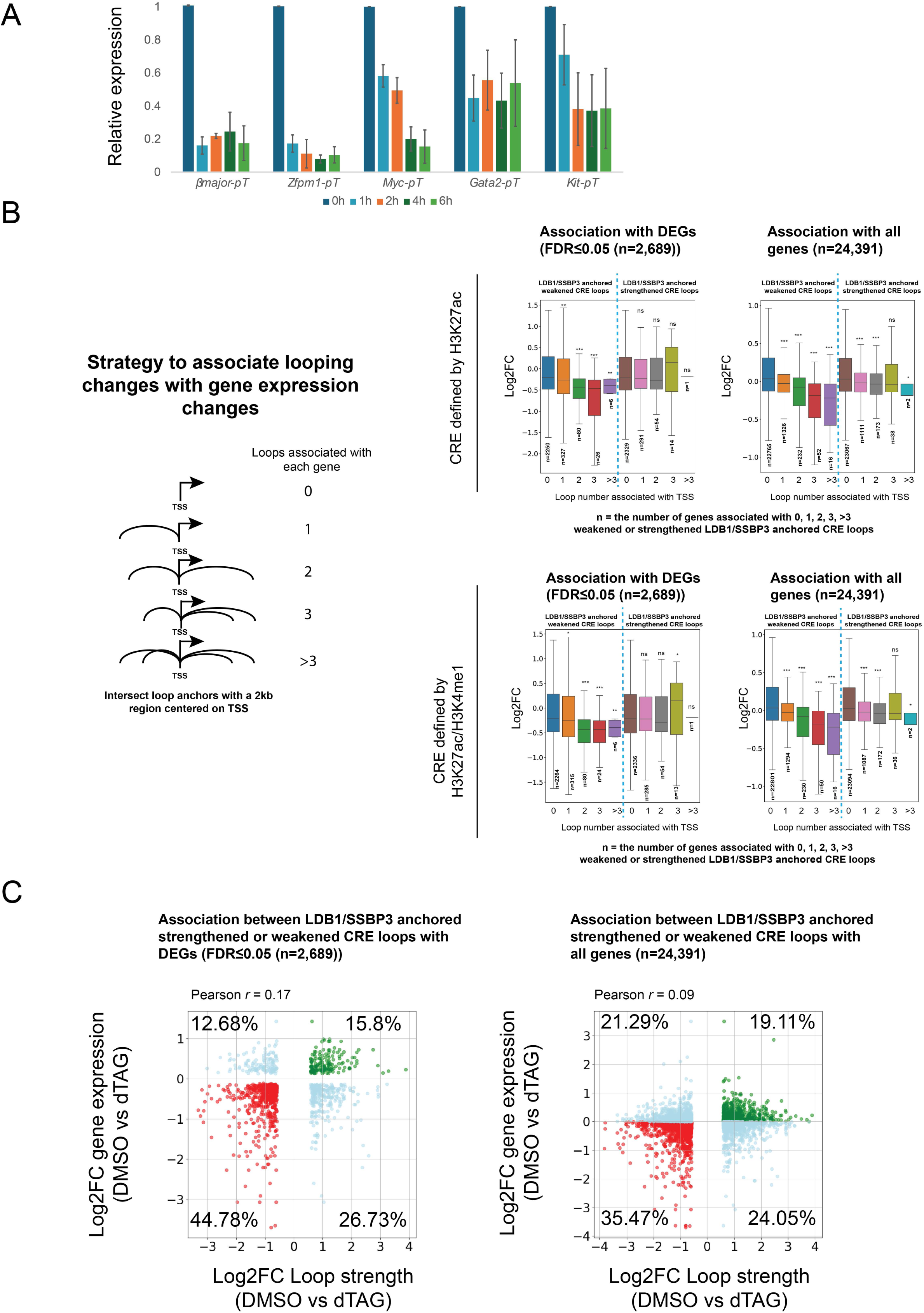
Downregulated genes after SSBP3 depletion are associated with weakened CRE loops, related to Figure 4. (A) RT-qPCR (n=3) to measure the primary transcripts of the indicated genes after a time series of 500 nM dTAG^V^-1 treatment. (B) Left: diagram showing the strategy to associate looping changes with gene transcription changes. Middle: Box plots showing LDB1/SSBP3 anchored weakened CRE loops are associated with downregulated gene, while LDB1/SSBP3 anchored strengthened CRE loops are not associated with upregulated gene. Differentially expressed genes (DEGs) with FDR≤0.05 (n=2,689) were used for association analysis. CRE is either defined by H3K27ac or by H3K27ac/H3K4me1. Right: Box plots showing LDB1/SSBP3 anchored weakened CRE loops are associated with downregulated gene, while LDB1/SSBP3 anchored strengthened CRE loops are not associated with upregulated gene. All genes (n=24,391) were used for association analysis. CRE is either defined by H3K27ac or by H3K27ac/H3K4me1. The box represents the interquartile range (IQR), with the line inside indicating the median. Whiskers extend to the smallest and largest values within 1.5 times the IQR. Statistical significance was assessed using the two-sided Mann-Whitney U test. * *p*≤0.05, ** *p*≤0.01, *** *p*≤0.001. (C) Scatter plot showing association between Log2FC of gene expression and Log2FC of loop strength. Left: Differentially expressed genes (DEGs) with FDR≤0.05 (n=2,689) were used for association analysis; Right: All genes (n=24,391) were used for association analysis.

**Figure S11.**
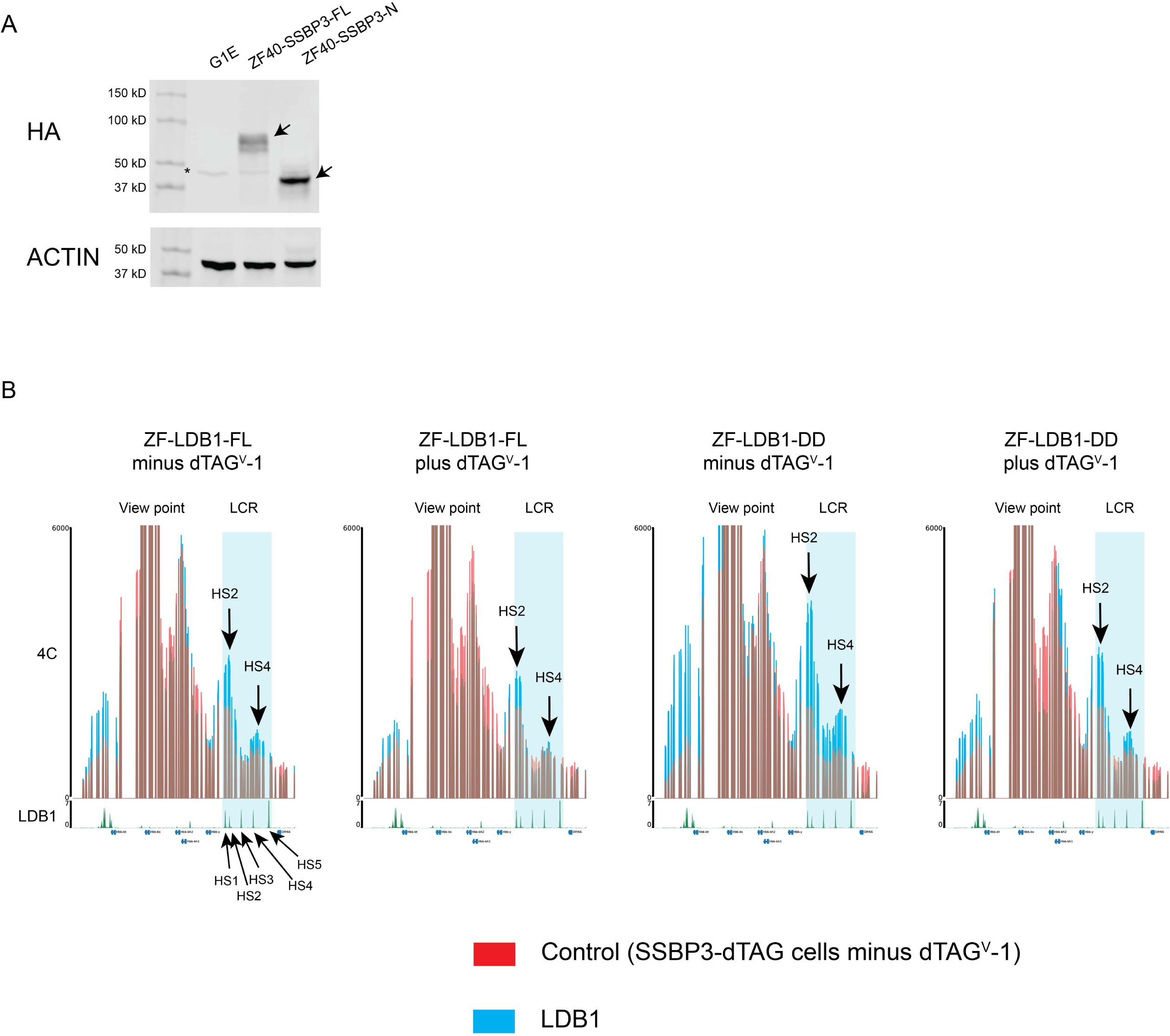
LDB1-meidated E-P interaction at β-Gloin locus requires SSBP3, related to Figure 5. (A) HA WB showing ZF-SSBP3-FL and ZF-SSBP3-N were expressed. ACTIN was used as a loading control for the inputs. WB was performed at least three times and representative result was shown. (B) 4C-seq results showing transfection of ZF-LDB1-FL (n=4) and ZF-LDB1-DD (n=3) into SSBP3-dTAG cells increased interaction between β*-major* promoter and the LCR enhancer. However, SSBP3 depletion decreased interaction between β*-major* promoter and the LCR enhancer. β*-major* promoter served as the viewpoint. The LCR enhancer was highlighted in cyan. LDB1 ChIP-seq track in undifferentiated G1E-ER4 was added below each 4C track.

